# Inferring and perturbing cell fate regulomes in human cerebral organoids

**DOI:** 10.1101/2021.08.24.457460

**Authors:** Jonas S. Fleck, Sophie M.J. Jansen, Damian Wollny, Makiko Seimiya, Fides Zenk, Malgorzata Santel, Zhisong He, J. Gray Camp, Barbara Treutlein

## Abstract

Self-organizing cerebral organoids grown from pluripotent stem cells combined with single-cell genomic technologies provide opportunities to explore gene regulatory networks (GRNs) underlying human brain development. Here we acquire single-cell transcriptome and accessible chromatin profiling data over a dense time course covering multiple phases of organoid development including neuroepithelial formation, patterning, brain regionalization, and neurogenesis. We identify temporally dynamic and brain region-specific regulatory regions, and cell interaction analysis reveals emergent patterning centers associated with regionalization. We develop Pando, a flexible linear model-based framework that incorporates multi-omic data and transcription binding site predictions to infer a global GRN describing organoid development. We use pooled genetic perturbation with single-cell transcriptome readout to assess transcription factor requirement for cell fate and state regulation in organoid. We find that certain factors regulate the abundance of cell fates, whereas other factors impact neuronal cell states after differentiation. We show that the zinc finger protein GLI3 is required for cortical fate establishment in humans, recapitulating previous work performed in mammalian model systems. We measure transcriptome and chromatin accessibility in normal or GLI3-perturbed cells and identify a regulome central to the dorsoventral telencephalic fate decision. This regulome suggests that Notch effectors HES4/5 are direct GLI3 targets, which together coordinate cortex and ganglionic eminence diversification. Altogether, we provide a framework for how multi-brain region model systems and single-cell technologies can be leveraged to reconstruct human brain developmental biology.

## Main text

The ability to generate complex brain-like tissue in controlled culture environments from human stem cells offers great promise to understand mechanisms underlying human brain development. Cerebral or brain organoids develop from pluripotent stem cells (PSCs) into a three-dimensional neuroepithelium that self-patterns, regionalizes, and ultimately forms neurons of the different brain regions (1–3). The fate and state of each cell is orchestrated through complex circuits of transcription factors (TFs) converging at regulatory elements to enable precise control of gene expression. Single-cell sequencing approaches allow the profiling of gene expression and chromatin accessibility in individual cells, which opens up new opportunities to survey the set of regulatory control features in any given cell fate or state (so-called regulomes). Direct comparisons between organoids and primary counterparts in mouse and human have quantified a remarkable similarity between the neural progenitor and neuronal transcriptome profiles (4–6). Cerebral organoids have been used to successfully model microcephaly (2), periventricular heterotopia (7), autism (8) and other neurodevelopmental disorders (9, 10) that may have differential effects on the various human brain regions. However, there is no detailed description of the gene regulatory networks (GRNs) that coordinate early human brain development.

Research in model systems have identified core signaling factors and gene regulatory programs that orchestrate brain region formation in vertebrates. Initially, extrinsic signals establish an anterior-posterior axis, which triggers additional localized gradients downstream to segment the neural tube into distinct brain regions. Combinatorial activities of morphogens including SHH, WNTs, BMPs, FGFs, NOTCH, Neuregulins and R-spondins converge on transcription factors to execute regionalization. The Sonic hedgehog (Shh) signalling pathway is a prominent component of this system that coordinates dorsoventral telencephalon specification. GLI3 belongs to the C2H2-type zinc finger proteins subclass of the Gli family, and is a terminal effector in the SHH pathway. In mice, loss of function mutations in GLI3 result in failure of the cortex to form, and expansion of ventral telencephalon neuronal identities into dorsal locations within the developing brain (11, 12). Conditional knock-out experiments have shown that GLI3 also plays an active role in regulating the onset of cortical neurogenesis by controlling cell cycle dynamics (13). Much of what is known about SHH, GLI3, and other pathways regulating brain morphogenesis have been explored in zebrafish and mouse model systems, and it remains unclear how human brain development has diverged from our mammalian ancestors. In humans, mutations in GLI3 have been associated with Greig cephalopolysyndactyly syndrome and Pallister Hall syndrome, in which patients have variable presentations of brain malformations depending on the particular mutations (14).

New single-cell genomic methods enable high-throughput and quantitative analysis of single-cell transcriptomes and accessible chromatin profiles. These features can also be measured within an individual cell in a multi-omic measurement, providing insight into gene expression and regulation in the same cell state. Furthermore, CRISPR-Cas gene editing coupled with single-cell transcriptome readouts (15–17) allows pooled genetic perturbation experiments in vivo (18). These strategies and vector systems, combined with functionalization of human PSCs with inducible Cas9 systems, provide an opportunity to perturb gene function in cerebral organoids, and systematically assess the effects across human brain regions.

Here we have used a multimodal approach to explore human brain regionalization. We first build a regulome from single-cell transcriptome and chromatin profiling data across a cerebral organoid developmental time course. We then perturb this regulome using multiplexed in organoid perturbation experiments, and identify effects on regional fate decisions as well as effects on cell states after fate acquisition. We show that multi-ome analysis of a critical period of brain region formation in isogenic control and GLI3 knock-out organoids distinguishes direct and indirect targets of this transcription regulator and reveals regulatory disruption of telencephalon development. Altogether, we establish a regulome-perspective to understand and explore early human brain development.

### Single-cell multiomic reconstruction of cerebral organoid development

To explore mechanisms underlying human brain development, we generated single-cell transcriptome and single-cell accessible chromatin profiling data over a time course of cerebral organoid development (Fig. 1A, Fig. S1A and data S1). The dataset incorporates 11 time points from four human PSC lines covering two months of development spanning embryoid body (EB) formation, neuroectoderm induction, neuroepithelialization, neural progenitor patterning, and neurogenesis. At each time point, tissues from the four lines were dissociated separately and combined into one single-cell suspension for both scRNA-seq and scATAC-seq pipelines (10X Genomics). The sequencing data was demultiplexed using single nucleotide variants (SNVs) specific to each individual and the two modalities for each line and time point were integrated using canonical correlation analysis (CCA) (19) (fig. S1, B to D). We constructed ‘multi-omic metacells’ containing information on both transcriptome and chromatin accessibility using minimum-cost, maximum-flow (MCMF) bipartite matching (20) within the CCA space (Fig. 1A). The metacells were integrated using cluster similarity spectrum (CSS) (21), and the integrated data was visualized using a Uniform Manifold Approximation and Projection (UMAP) embedding (fig. S1, E and F, and data S2). This revealed a relatively continuous distribution of cell states through the entire time course (Fig. 1, B to D, and fig. S1, G and H). Organoid development proceeds from pluripotency (e.g. POU5F1) via a neural progenitor cell state (e.g. PAX6, VIM1) to progenitor and neuron cell states of the dorsal telencephalon (e.g. EMX1, NEUROD6), the ventral telencephalon (e.g. DLX5, ISL1, GAD1) as well as of non-telencephalic regions (e.g. TCF7L2, LHX9), with cells from the different lines largely intermixed. The high-dimensionality of the data could be used to identify marker genes and gene regulatory regions for the different cell states, as well as promoter accessibility and binding motif enrichment for stage-specific transcription factors (TFs) (Fig. 1, D and E, and data S3). We observed a pseudotemporal cascade of chromatin accessibility changes over the developmental time course associated with genes involved in stem cell maintenance, neural tube patterning, morphogenesis, neural precursor proliferation, neuron fate specification, and other relevant biological processes (Fig. 1, F and G, fig. S1I, and data S4). Generally, the genome showed highest accessibility in early states associated with pluripotency and the transition to neuroepithelium, whereas it progressively restricted in later stages of organoid development.

**Fig. 1.**
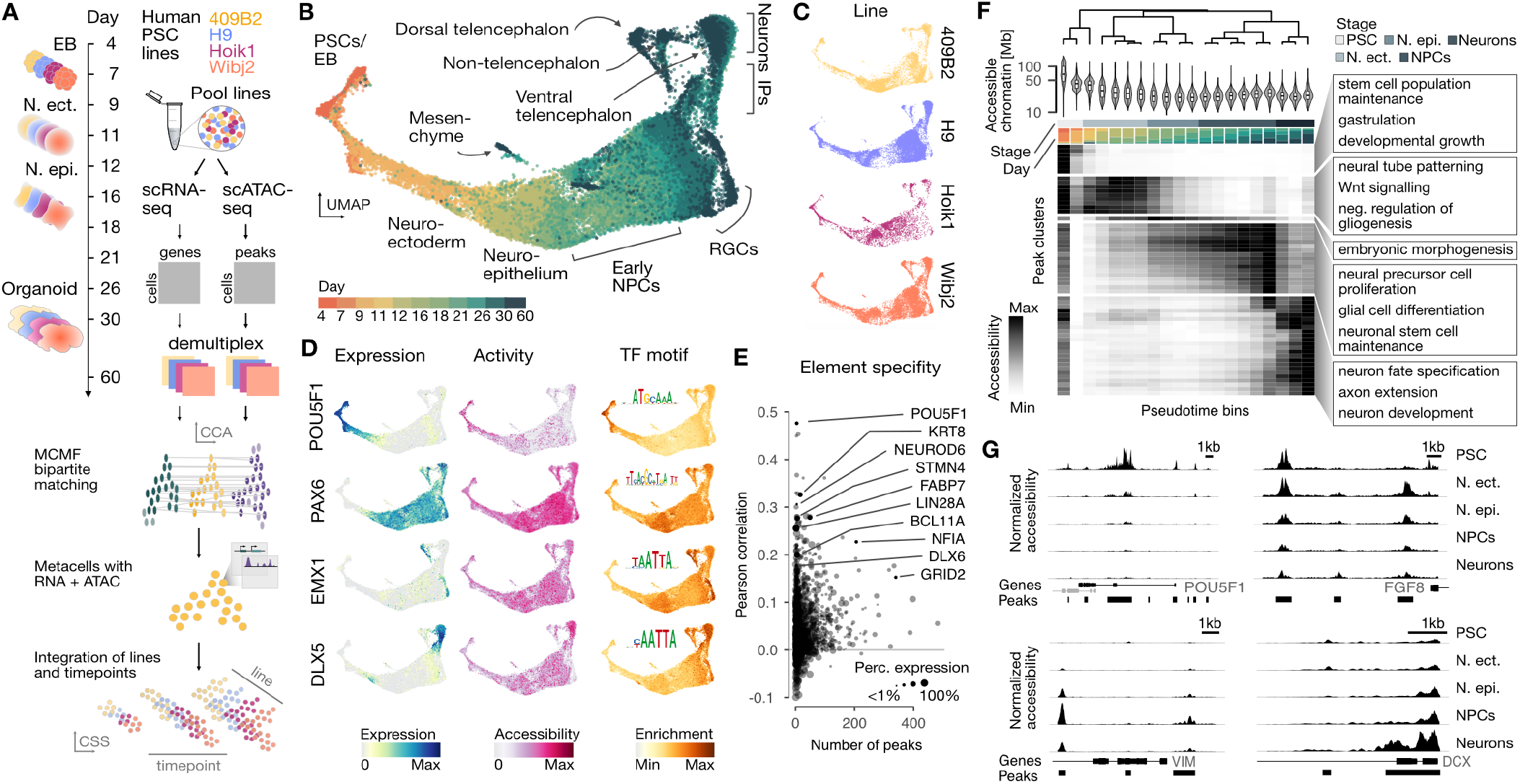
Gene expression and chromatin accessibility dynamics during early human cerebral organoid development. (A) Schematic of the experimental design and data integration strategy. Organoids from four different iPSC lines were dissociated for scRNA-seq and scATAC-seq at timepoints spanning the critical patterning window. Data was demultiplexed using SNV information and modalities can be integrated using minimum-cost, maximum-flow (MCMF) bipartite matching in CCA space. The resulting metacells with RNA and ATAC components are integrated using CSS (21). (B) UMAP embedding of the full integrated dataset. (C) Distribution of iPSC lines on the UMAP embedding. (D) Gene expression (left), gene accessibility (middle) and binding motif enrichment (right) for stage-specific transcription factors. (E) Scatter plot showing the Pearson correlation between gene accessibility and gene expression versus the number of peaks in gene body and promoter regions. (F) Hierarchical clustering of pseudotemporal bins revealing regulatory transitions. Violin plots show the total length of accessible chromatin per bin, top side bars show stage (grey scale) and proportion of cells at each time point (stacked plot) per bin. Heatmap shows accessibility of stage-specific peak clusters for each pseudotime bin. Representative GREAT enrichments are shown for each stage. (G) Examples of loci that have differential access during cerebral organoid development from pluripotency.

We next wanted to reconstruct how regional heterogeneity emerges during cerebral organoid development. Towards this aim, we sub-clustered early portions of the trajectory and identified molecular heterogeneity (Fig. 2A and fig. S2). In the initial stages (day 7-9), we observed a predominant neuroectodermal population (VIM, SIX3, CDH2, SOX3, HES5) and a minor population of cells expressing non-neural ectoderm markers (DLX5, TFAP2A) (22) (17, 23) (fig. S2, A to C). In the neuroepithelium (day 12-18), expression gradients emerged associated with patterning centers including the antihem (PTX3, DLX2, GSX2), floor plate (RAX, SIX3, SIX6) and roof plate (LMX1A, MSX1, DMRT3, WNT3A) (fig. S2, D and E). We compared these organoid patterns to a scRNA-seq map of the developing mouse brain (24), and found similarity to populations that organize dorsoventral and anterior-posterior gradients (Fig. 2B and fig. S2, F and G). Over time, states associated with these centers further diverged into neural progenitor cells (NPCs) expressing either telencephalic and non-telencephalic markers, followed by a second divergence into dorsal and ventral telencephalic NPCs (Fig. 2C and fig. S2, H to K). Intriguingly, we identified clusters within the early neuroepithelium that have a significant enrichment of receptor-ligand interactions with multiple other clusters, suggesting the presence of organizing centers within the developing organoid (fig. S3). These data extend previous work showing that patterning centers emerge in the neuroepithelium, which coordinate to regionalize the developing organoid (25).

**Fig. 2.**
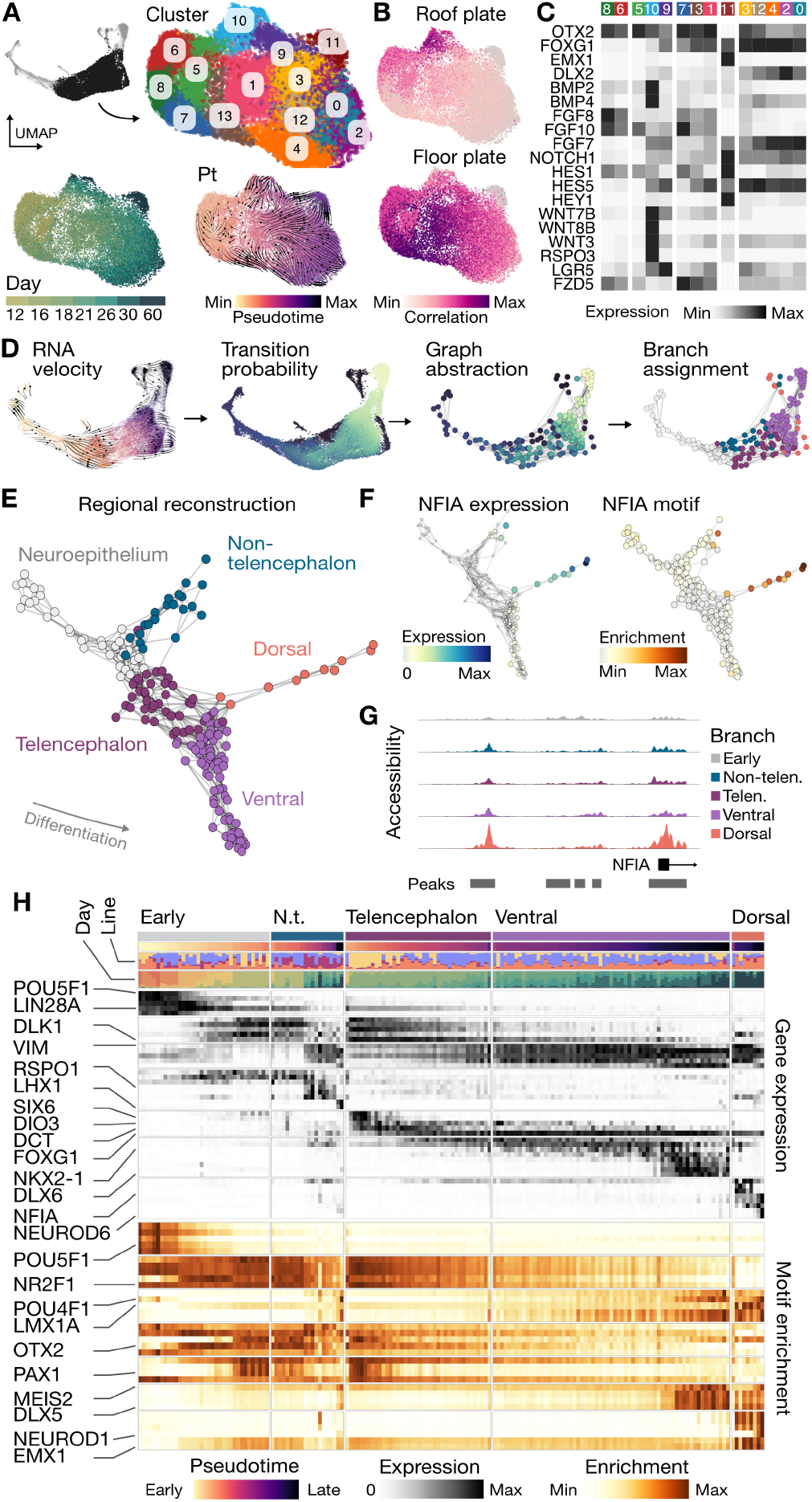
Resolving cerebral organoid patterning and regional fate trajectories. (A) UMAP embedding of subsetted neural progenitor cells colored by Louvain clusters, time point and velocity pseudotime. (B) UMAP embedding colored by the correlation to developing mouse brain organizers. (C) Heatmap showing the normalized expression of cluster marker genes related to brain patterning. (D) Schematic of branch-inference strategy. High-resolution clusters were assigned to branches based on terminal fate transition probabilities calculated based on RNA velocity. (E) Branch visualization in a force-directed layout, with circles representing high-resolution clusters with both RNA and access features colored by assignment (Neuroepithelium, grey; Non-telencephalon progenitors, teal; Telencephalon progenitors, plum; Dorsal telencephalon, orange; Ventral telencephalon, purple). (F) Graph representation of regional branches colored by NFIA gene expression (left) and binding motif enrichment (right) (G) Signal tracks showing normalized accessibility at the transcription start site of NFIA in the different branches. (H) Heatmap showing stage- and branch-specific gene expression and motif enrichment.

From the regionalized NPCs, we next sought to reconstruct the neurogenic differentiation trajectories for each brain region. We used RNA velocity (26, 27) and CellRank (28) to generate a terminal fate transition probability matrix based on transcriptomes, which we used to construct a differentiation graph of high-resolution metacell clusters and assign branch identities (Fig. 2D, and figs. S2J and S4, A to E). The graph, presented by a force-directed layout, reveals an early bifurcation into anterior telencephalic and posterior non-telencephalic cell states and later branching of telencephalic progenitors into dorsal excitatory and ventral inhibitory neuronal trajectories, respectively (Fig. 2E). Transcriptional and regulatory dynamics can be explored along each neurogenic trajectory, revealing regional specificity of gene expression and chromatin accessibility (Fig. 2, F to H, and fig. S4, F and G). As examples, we highlighted NFIA as regulator of neurogenesis in the dorsal telencephalon, and also identified a telencephalic progenitor state prior to dorsoventral divergence marked by the expression of DCT, DIO3 and SIX6, and characterized by transient accessible chromatin regions. Altogether, this data provides a multi-omic developmental atlas spanning the course of brain organoid regionalization and neurogenesis.

### A gene regulatory network view of human cerebral organoid formation

Next we wanted to harness our multi-omic single-cell data to infer the gene regulatory network (GRN) underlying human cerebral organoid development. We developed an algorithm called Pando (Fig. 3A, see Methods), which incorporates scATAC-seq data to identify conserved non-exonic regions and candidate cis-regulatory elements (29) (cCREs) that are accessible across the organoid time course (candidate regions, fig. S5, A and B). TF binding sites are predicted for each candidate region (fig. S5, C to E), and the relationship between TF expression and expression of the potential target genes with binding sites within nearby regions is inferred using a linear model (fig. S5F). As a consequence, Pando jointly infers sets of positively or negatively regulated target genes (gene modules) as well as regulatory genomic regions (regulatory modules) for each TF (fig. S5, G and H). We visualized the GRN using a UMAP embedding, which revealed groups of TFs that are involved in different phases of cerebral organoid development, broadly representing the pseudotemporal order of cell state transitions (Fig. 3B and fig. S5I). A series of TFs tracked transitions from pluripotency (e.g. POU5F1, LIN28A) to neuroepithelium induction (e.g. SOX2, HES1), with additional module neighborhoods linked to brain regional NPC specification and neuron differentiation (Fig. 3C). Nodes associated with initializing (pluripotency) and terminal states (regionalized neurons) had a high degree of centrality, reflecting the high number of correlated expressed genes for these states. Globally, this GRN shows that regulatory region accessibility and TF expression track with stages of organoid development and segregate during brain regionalization.

**Fig. 3.**
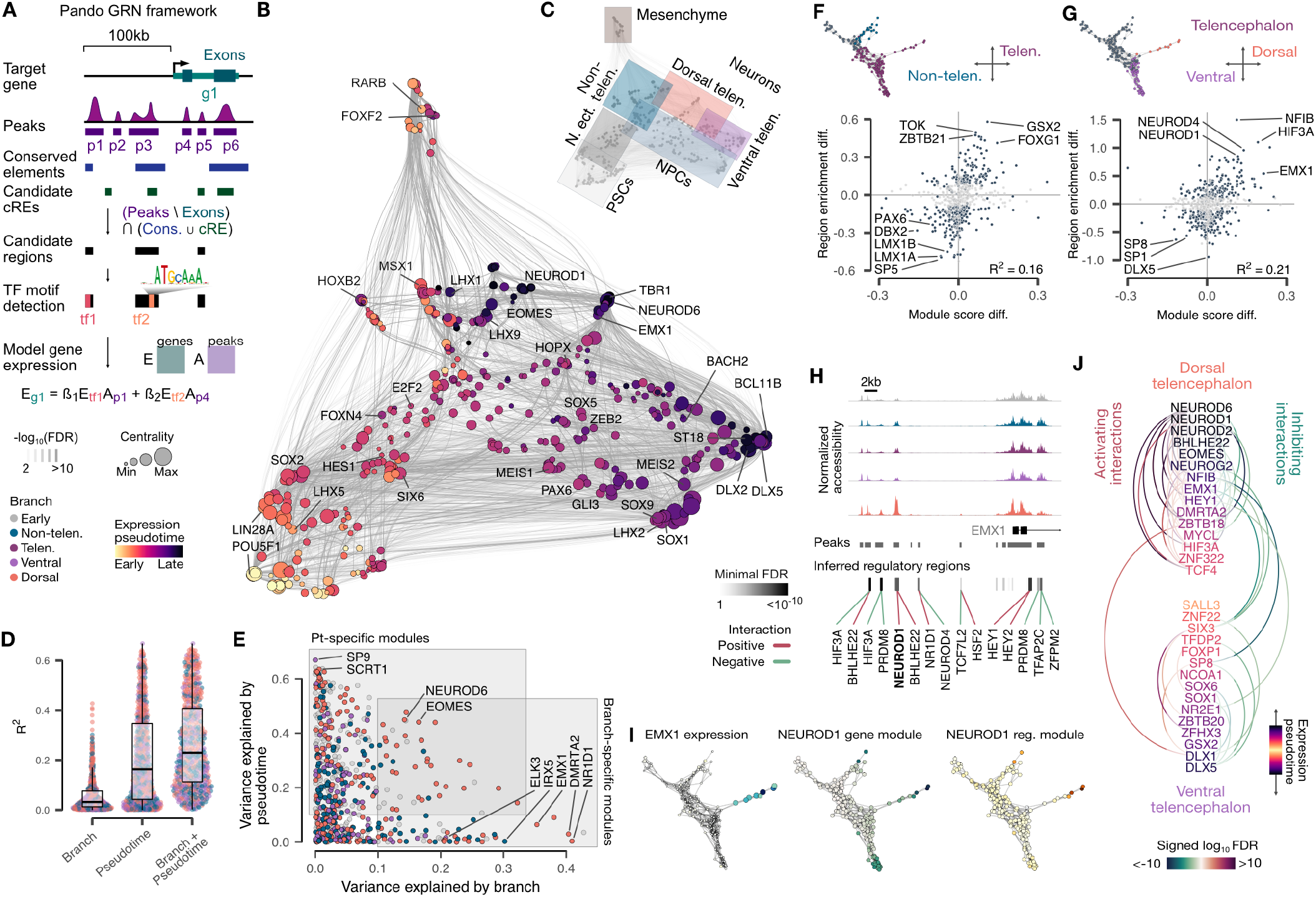
Gene regulatory network underlying human cerebral organoid formation. (A) Schematic of the Pando GRN-inference framework. Candidate regions are identified through intersection of non-exonic accessible peaks with cis-Regulatory Elements (cREs) or conserved elements. Predicted TFs are selected for each candidate region through binding motif matching. A linear model explains the relationship between TF-binding site pairs and expression of target genes. (B) UMAP embedding of the inferred gene modules based on co-expression and inferred interaction strength between TFs. Color and size represent expression weighted pseudotime of TF regulator and pagerank centrality of each module. (C) UMAP embedding shaded by module features. (D) Variation of module activity explained by branch, pseudotime, or branch and pseudotime. Circles represent individual modules. (E) Branch and pseudotime specific modules in the GRN. Colors represent the branch with highest average module activity. Modules where the TF motif was not experimentally validated are shown in grey. (F-G) Branch specific gene and regulatory modules for the telencephalon / non-telencephalon split (F) for the ventral / dorsal telencephalon split (G). (H) Signal tracks showing normalized accessibility at the transcription start site of EMX1 in the different branches and inferred regulatory regions for various transcription factors. Line color represents the sign of the interaction and box color (greyscale) represents the FDR of the most significant interaction for this region. (I) Graph representation of regional branches showing the expression of EMX1 as well as gene and regulatory module activity for NEUROD6. (J) Interactions between dorsal and ventral telencephalon-specific transcription factors ordered by expression pseudotime.

We next sought to better understand neuronal trajectory relationships within the GRN. We analyzed the variance explained by brain region (branch) and by general developmental pseudotime for all TF modules in the GRN (Fig. 3, D and E). We found that certain TF modules (e.g. SP9, SCRT1) were highly pseudotime dependent, and were correlated with neurodevelopmental dynamics in multiple brain regions. In contrast, other TF modules were specific to a brain region but were not dynamic along the neurogenic pseudotemporal trajectory (e.g. EMX1, NR1D1 in dorsal telencephalon; IRX5 in non-telencephalon). Another set of TFs had strong variance explained by both pseudotime and branch and these TF modules represent maturation state- and fate-specific features within the developing organoid (e.g. NEUROD6, dorsal telencephalon neurons; EOMES in dorsal telencephalon intermediate progenitors). Indeed, we could use the GRN to reveal gene regulatory programs that diverge upon specification of progenitors and differentiation of neurons into organoid brain regional fates (Fig. 3, F and G). For instance, gene and regulatory modules for NFIB, NEUROD1 and EMX1 are specific to the dorsal telencephalon and characterize the regulatory changes tracking the specification of this developmental trajectory. As a representative example, we show the EMX1 locus and highlight predicted TF regulators such as NEUROD1 within accessible chromatin regions (Fig. 3, H and I). More broadly, we could infer TF modules that distinguish dorsoventral neuron fate specification and differentiation in the telencephalon (Fig. 3J). Altogether, these analyses provide a rich resource for future work to understand the gene regulatory programs controlling human brain regionalization and cell programming.

### *In organoid* single-cell genomic perturbation screen

To begin to understand the mechanisms regulating cell fate and state during human brain development, we employed a pooled perturbation screen (17) in mosaic organoids (Fig. 4A). We designed gRNAs targeting 20 TFs expressed in both the organoid and primary developing human cortex (4) (Fig. 4B and fig. S6, A and B), and generated a pooled lentiviral library. We infected induced pluripotent stem cells (iPSCs) harboring an inducible Cas9 cassette with the lentiviral gRNA library, and sorted and expanded vector positive iPSCs based on fluorescence (fig. S6C). We induced Cas9 expression in the infected iPSCs, and generated mosaic cerebral organoids containing a multitude of wild-type (WT) and knock-out (KO) genotypes (fig. S6B and C). Fluorescence was maintained throughout organoid development, and bulk amplicon sequencing revealed relatively homogenous detection of the gRNAs (figs. S6D and S7A). At day 60, we sequenced single-cell transcriptomes and guide cDNA amplicons and recovered 22,449 cells with an assigned gRNA (Fig. 4C, and fig. S7, B to D). Each gRNA for all 20 targets was detected with an average of 1 gRNA detected per cell (fig. S7, D to F). We generated a UMAP embedding, analyzed cell type heterogeneity, and annotated NPCs, intermediate progenitors, and neurons in the dorsal telencephalon, the ventral telencephalon, as well as in non-telencephalic developing brain regions (Fig. 4C, and fig. S7, G to I). To determine the effect of TF perturbation on cell fate, we first tested the association of gRNA detection with cell type abundance. We hierarchically clustered all Louvain clusters on the basis of gRNA abundance and observed grouping based on brain region identity (fig. S7J). This showed that different brain regions exhibited unique gRNA compositions suggesting region specific effects of TF KOs. Next, we stratified the detected gRNAs using a log odds ratio (p-value based on a Cochran–Mantel–Haenszel test) and the consistency of the effect across organoids and gRNAs (see Methods), and visualized these metrics across the different brain region fates in a ‘lollipop plot’ (Fig. 4D, and data S5). Based on these metrics, we found that for 8 TFs there was a consistent enrichment (support from at least 2 gRNAs) in the ventral telencephalon branch with corresponding depletion in the other regions, including the cortex (Fig. 4E). Among these, we highlight TBR1 and GLI3, two strong effect genes with consistent depletion in the dorsal telencephalon and enrichment in the ventral telencephalon (Fig. 4F). Both genes are known regulators of mouse cortical development (30, 31), and are associated with neurodevelopmental disorders in humans (14, 32).

**Fig. 4.**
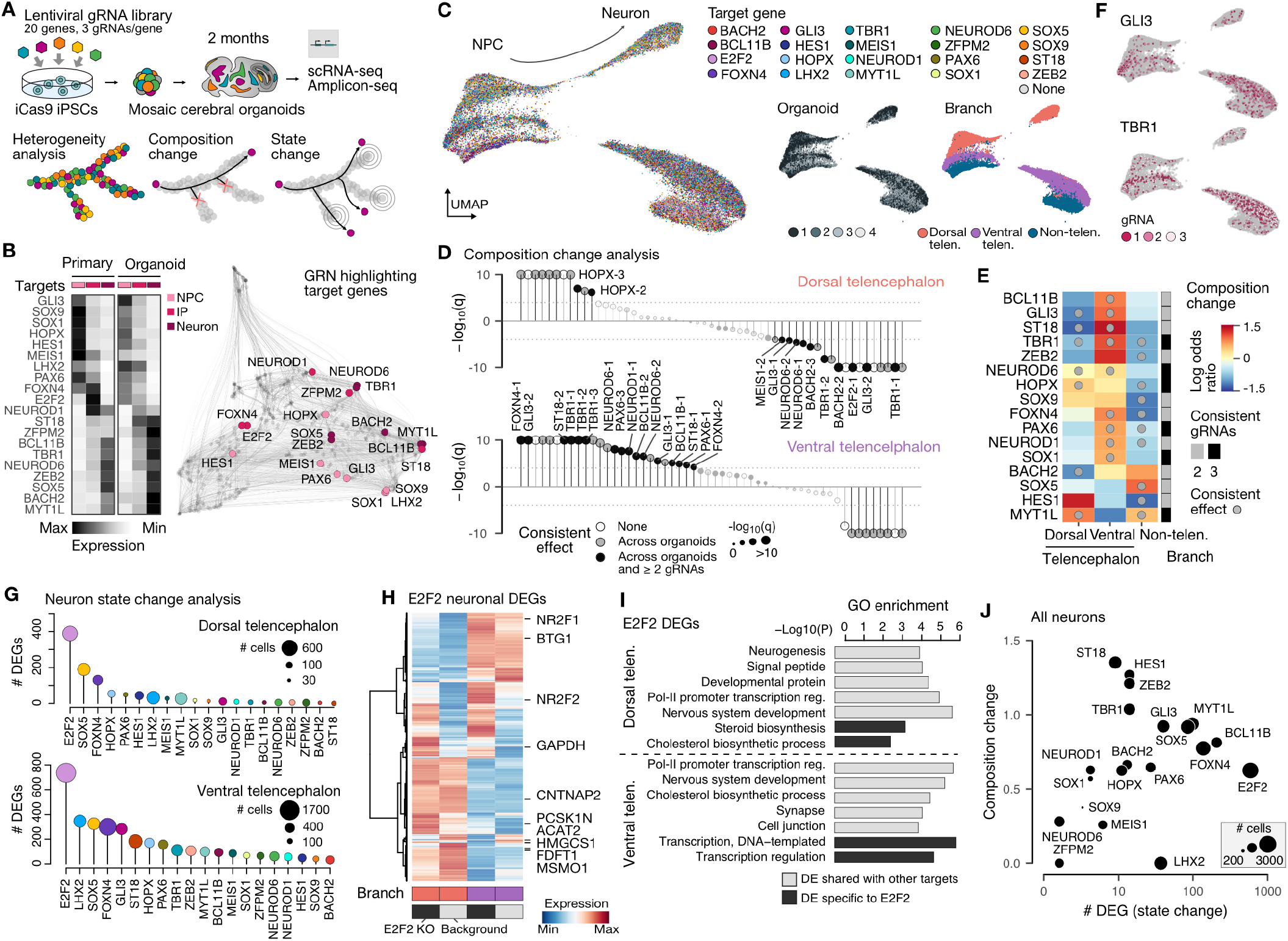
*In organoid* single-cell genomic perturbation of cortical transcription factors. (A) Schematic of single-cell perturbation experiment using the CROP-seq method and data analysis. (B) Heatmap showing average expression of targeted genes in neural progenitor cells (NPC), intermediate progenitors (IP) and neurons of the primary and organoid cortex. Targeted TF nodes are highlighted on the GRN. (C) UMAP embedding with cells colored based on the detected gRNA, organoid sample and branch assignment. (D) Lollipop plot showing the impact of each gRNA on cell type abundance in dorsal and ventral telencephalic neurons. (E) Heatmap showing effect of gRNAs on abundance of regional branches. Sidebar shows the number of gRNAs that were consistent and circles represent consistent and statistically significant (FDR<0.01) effects on composition. (F) UMAP embedding colored by detected gRNAs for selected genes that had strong effect on fate regulation. (G) Lollipop plots showing number of differentially expressed genes (DEG) for targeted genes in the dorsal and ventral telencephalic neurons. (H) Heatmap of E2F2 DEGs in the dorsal and ventral neuron. (I) Examples of functional enrichment for E2F2 DEGs in dorsal and ventral neurons. (J) Differential gene expression analysis was performed to identify potential effects on cell state. Plot shows the effect of cell composition change and the number of differentially expressed genes (DEGs).

We next performed differential gene expression analysis to determine perturbation effects on gene expression in dorsal and ventral telencephalic neurons (Fig. 4G, and data S6). Interestingly, for both neuron types we detected the most differentially expressed genes (DEGs) for E2F2, a crucial cell cycle regulator (33) that has enriched expression in intermediate progenitors. These E2F2-related changes in the two neuron types were distinct but correlated (Fig. 4H and fig. S8, A and B), and some of the DEGs were specific to E2F2 when compared to effects from other targets (fig. S8, C and D). Functional enrichment analysis suggests that E2F2 DEGs are significantly related to neuron development and neuron-related function (Fig. 4I, fig. S8E, and data S7). This suggests that misregulation of cell cycle exit during the transition from a progenitor to neuron state has a large effect on the neuronal transcriptome state. For all targeted genes, we compared the number of DEGs with the extent of composition changes and did not observe substantial correlation, implying independent mechanisms involved in the two types of changes (Fig. 4J). Altogether, our data provides one of the first implementations of an *in organoid* multiplexed perturbation experiment to systematically understand the effect of putative gene KO on human brain cell fate and state development.

### GLI3 regulates the dorsoventral fate decision in the human telencephalon

Mosaic perturbations suggested that GLI3 is involved in dorsoventral neuronal fate specification in the human telencephalon. To confirm this result, and to explore the underlying developmental mechanisms, we used CRISPR/Cas9 gene editing to generate two isogenic GLI3 KO iPSCs and control cells (WT) that went through the editing process (Fig. 5A, and fig. S9, A to D). We generated KO and WT cerebral organoids and confirmed that the GLI3 protein is not detected in the KO organoids (fig. S9, C and E, and data S8). We performed scRNA-seq on KO and WT organoids at day 45, a time point of early neurogenesis, and analyzed cell heterogeneity (Fig. 5, B and C). We observed a striking effect in that the isogenic KO cells were fully depleted in the dorsal telencephalon, with a strong enrichment in the ventral telencephalon (Fig. 5D), consistent with the mosaic perturbation experiment. We analyzed the effect of GLI3 KO on ventral telencephalic cell states using differential expression analysis, and found strong correspondence in the results from the isogenic and mosaic perturbation experiments (Fig. 5E, and data S9). Interestingly, we found that in both cases the TF MEIS2, which is also a marker of lateral/caudal ganglionic eminence (LGE/CGE) relative to medial ganglionic eminence (MGE) was strongly down-regulated in GLI3 KO conditions (Fig. 5E). Further analysis on the ventral telencephalic neuron heterogeneity identified distinct LGE/CGE-like and MGE-like neuronal populations with GLI3 KO cells strongly enriched in MGE neurons (Fig. 5F, and fig. S9F). Interestingly, we observed expression alterations in GLI3 KO LGE-like neurons compared to the WT LGE state with genes involved in dorsoventral patterning (PAX6, MEIS2, DLK1) being differentially expressed (fig. S9G). These data confirm that GLI3 is required for cortical fate establishment in humans, and its absence impacts development of the ventral telencephalon by promoting the emergence of MGE neurons consistent with a role as repressor of the MGE fate (34) and by altering the state of LGE neurons (Fig. 5G).

**Fig. 5.**
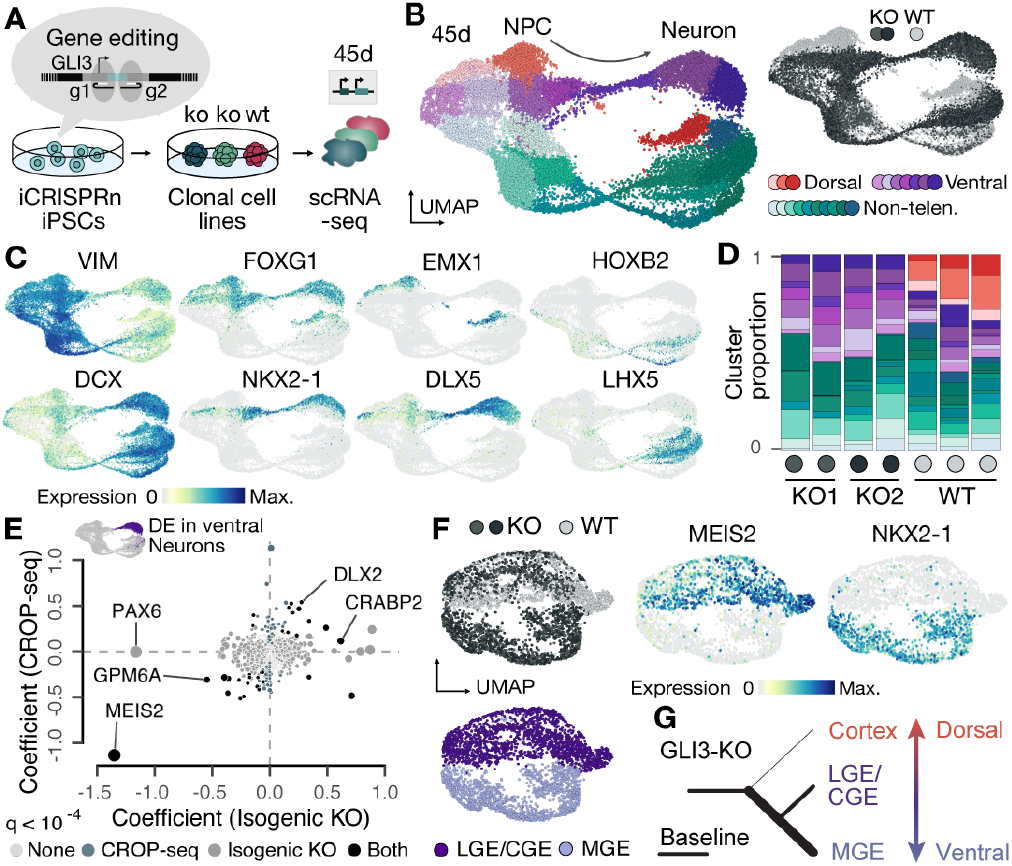
GLI3 regulates the dorsoventral telencephalic fate decision in humans. (A) Schematic of GLI3 loss of function experiment (knock-out, KO; wild-type, WT) using the iCRISPR nickase system. (B) UMAP embedding showing trajectories from neural progenitor cells (NPCs) to neurons colored by different clusters assigned to branches (dorsal, ventral, and non-telencephalon), with inset colored by genetic condition. (C) Feature plots colored by expression of cell type markers. (D) Stacked barplots showing distribution of cluster assignment per organoid for each condition, colored by cluster. (E) Differential expression (DE) in ventral telencephalic neurons for GLI3 isogenic KO and GLI3 CROP-seq data. X and y axes indicate coefficients of the linear model. Colors indicate whether the gene was significant (FDR<10^−4^) in CROP-seq, isogenic KO, or both. (F) Sub-clustering ventral telencephalic GLI3 KO neurons reveals medial ganglionic eminence (MGE) and lateral/caudal ganglionic eminence (LGE/CGE) populations. UMAPs colored by genetic condition, GE population, and marker expression level. (G) Schematic of observed effect of GLI3 loss of function on dorsoventral telencephalic fate decisions.

### GLI3 is a mediator between SHH, NOTCH and WNT signaling pathways

To identify gene regulatory mechanisms controlling dorsoventral fate decisions in the human telencephalon and to further illuminate the role of GLI3 in this process, we generated single-cell multiome data (10X genomics) measuring both transcriptome and chromatin accessibility from the same single cells derived from 3 week old WT and GLI3 KO organoids (Fig. 6, A and B, and fig. S10, A to C). This time point was chosen based on our previous analysis showing that cell states were present in the organoids that surround the dorsoventral bifurcation decision. The multiome data revealed strong transcriptomic differences between KO and WT particularly in the early telencephalic progenitor population (cluster 0, Fig. 6, B and C, fig. S10D, and data S10). Examining the expression of ligands, receptors, and target genes of several important signaling pathways suggested substantial changes on the cell-cell communication landscape in the GLI3 KO organoids, including the significant differential expression of Notch targets HES1 (up-regulated) and HES5 (down-regulated) in the telencephalic progenitor population (cluster 0 and 2, Figure 6D). Interestingly, promotion of dorsal telencephalic fate emergence by HES1 was observed in the CROP-seq experiment (Fig. 4E), in contrast to depletion of the dorsal telencephalic fate in the GLI3 KO. Some of the observed DEGs in NPCs containing HES1 gRNAs in the CROP-seq experiment were also observed in the GLI3 KO telencephalic progenitor population (SOX4, SOX11), further implying crosstalk between the pathways (fig. S10E). Differential ligand-receptor pairing analysis highlighted the same population of early telencephalic progenitors (cluster 0, 2, 6 and 12) for its strengthened role as a signaling source in the GLI3 KO organoids (fig. S10, F to K).

**Fig. 6.**
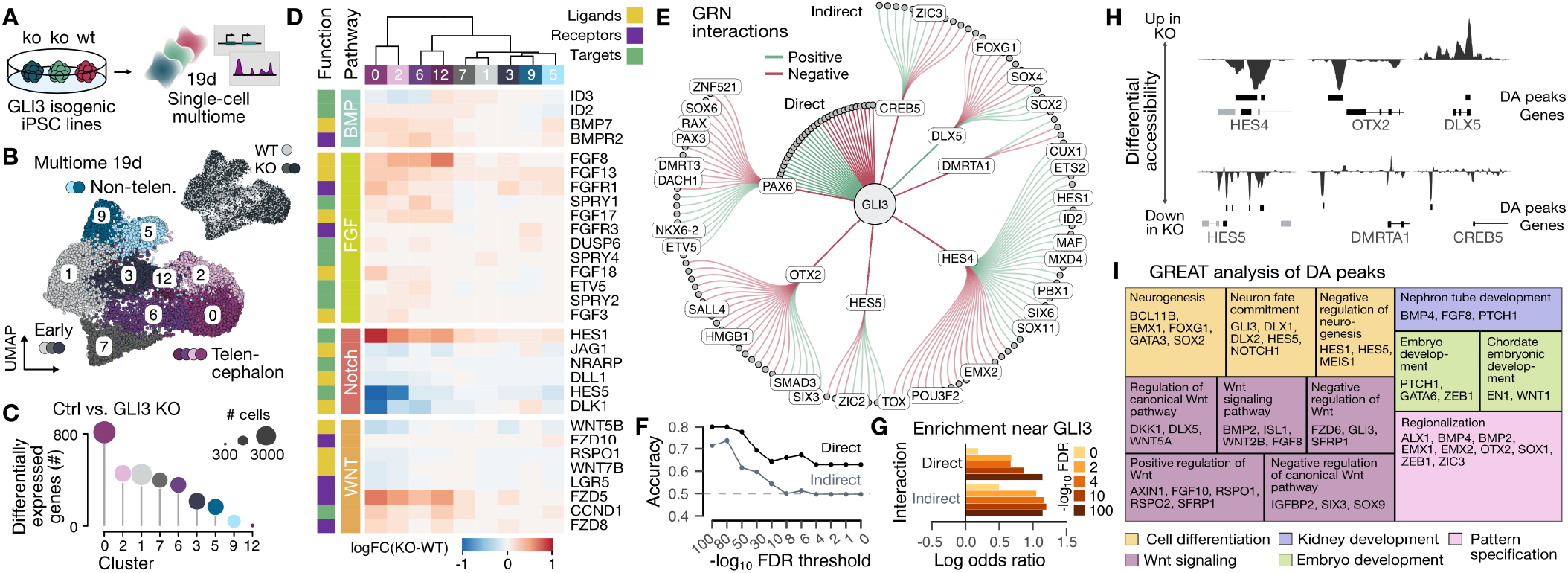
Telencephalic dysfunction of WNT and NOTCH regulomes in the absence of GLI3. (A) Schematic of the experiment where transcriptome and chromatin access was measured in the same cell at 19 days of cerebral organoid development. (B) UMAP embedding colored by cluster, and labeled by projected cell fate. Inset UMAP colored by genetic state. (C) Lollipop plot showing number of DEGs of control (Ctrl.) versus GLI3 KO cells in the different clusters. Circle color and size indicate cluster and number of cells, respectively. (D) Heatmap showing DE associated with various signaling pathways. Genes are classified as ligand (L), receptor (R) and target (T). (E) Subgraph of the GRN, showing direct and second-order indirect GLI3 targets whose inferred regulatory interaction is consistent with the observed DE. Circles are genes with all TFs being labelled. Edge interactions are colored based on positive (teal) or negative (maroon) regulation by the TF. (F) Accuracy of GRN predicted directionality of GLI3 effect for direct and indirect interactions at different false discovery rate (FDR) thresholds. (G) Enrichment of DE genes in the direct and indirect neighborhood of GLI3 in the GRN graph. (H) Signal tracks showing differentially accessible (DA) peaks in cluster 0 and 2 nearby genes that are differentially expressed between GLI3 KO and WT conditions. (I) GREAT enrichment analysis of DA peaks in cluster 0 and 2, with box area proportional to FDR and colored by ontology category. Representative genes near DA peaks are shown for each category.

To better understand how the observed changes were regulated by GLI3 loss of function, we used the global GRN to distinguish direct from indirect effects of GLI3 and to construct a mechanistic model of the perturbation signature in telencephalic progenitors (Fig. 6E, and fig. S10, L and M). We tested whether the effects observed in the KO were consistent with the previously inferred regulatory network and found that DEGs were highly enriched among direct and indirect targets of GLI3 (Fig. 6F). 76% of the DEGs were indirect targets of GLI3, but downstream of other TFs directly regulated by GLI3 such as HES4, HES5, OTX2 and PAX6. Moreover, for the most highly differential genes, the GRN could predict the directionality of the interaction with an accuracy of up to 80% for direct and up to 74% for indirect interactions (Fig. 6G). Interestingly, both the direct and indirect transcriptomic changes correlated positively with transition probabilities to ventral telencephalon and negatively with transition probabilities to dorsal telencephalon (fig. S10N), consistent with the disrupted cortical fate establishment and ventralization of the telencephalon in GLI3 KO organoids. We identified differentially accessible (DA) peaks between WT and GLI3 KO organoids, and found substantial KO-induced alterations in chromatin state in the genomic vicinity of DEGs (Fig. 6H, and data S11). GREAT enrichment analysis (35) revealed an association of differentially accessible regulatory regions with neuron differentiation, the WNT pathway and regionalization (Fig. 6I, and data S12). Altogether, this data suggests that GLI3 is a mediator between SHH, NOTCH and WNT signaling pathways.

## Discussion

The human brain has unique features that distinguish it from other species. It has been a major challenge to study the mechanisms that control brain development due to difficulty in obtaining tissue across a time window spanning the different stages of development, and the lack of methods to systematically manipulate gene function. Here we have integrated transcriptome, chromatin accessibility, and genetic perturbation datasets to provide insight into the mechanisms underlying human brain regionalization. In a broad sense, we find that the programs identified in mouse and other non-human model systems are well conserved in humans, and the extent that stem cell-derived cerebral tissues recapitulate these programs is remarkable. As a proof of principle, we focused on GLI3 as a well-studied transcription factor controlling dorsoventral fate specification in the rodent telencephalon. We find clear and striking evidence that this same transcriptional program is conserved in humans. More importantly, this data provides strong evidence that multi-region human cerebral organoids can be predictive model systems.

We established the Pando GRN inference framework which incorporates features of the regulatory genome that have not been previously utilized for global analysis of developmental programs. Pando combines transcriptome, chromatin accessibility, an expanded TF family motif reference, known cis-regulatory elements, and evolutionary conservation into a flexible, linear model-based framework. The package implements the full GRN inference strategy including candidate region selection, motif matching, model fitting and discovery of gene and regulatory modules. We have highlighted interesting aspects of the network, such as TF modules involved in the transition from pluripotency via neuroectoderm to a neuroepithelium, as well as the modules associated with regionalized brain states. The latter modules will be interesting to explore in experiments designed to program specific neuronal states, or to systematically perturb human organoids. We validated the critical role of GLI3 in dorsal telencephalic cell fate specification in humans, and further identified the contribution of GLI3 during specification of MGE and LGE/CGE neurons. The integration of the single-cell multiome data from GLI3 KO organoids and the global GRN allowed us to propose a model in which GLI3 becomes induced in early telencephalic NPCs through SHH signaling during neuroepithelial regionalization. GLI3 then regulates downstream targets, activating cortical fate acquisition and inhibiting the MGE induction program, through differential activity of Notch target genes HES5, HES4 and HES1. Notably, HES1 knockout leads to an enrichment of dorsal telencephalic cell states, providing further evidence for extensive crosstalk between the WNT and NOTCH pathways during human brain regionalization and neurogenesis. More broadly, our data reveals the extraordinary potential of multimodal single-cell genomic and organoid technologies to understand gene regulatory programs of human brain development.

There are advantages and limitations to the protocol that we used for *in organoid* genetic perturbations. On the positive side, inducible Cas9 iPSCs enable the temporal control of genetic manipulation. In this project, we perturbed genes at the iPSC stage, however KO can in principle be induced at any stage of organoid development. The vector system we used contained a single gRNA, and other approaches with cloning sites for 2 gRNAs may enhance KO efficiencies. Further modifications to the vector system that enable simultaneous genetic perturbation and lineage recording could be used to provide additional modalities to the measurements (36). We utilized the propensity of different human iPSC lines to give rise to a spectrum of human brain regions using the same cerebral organoid protocol in order to identify patterning and region-specific regulomes. However, it is a major challenge to use the current multi-region brain organoid protocols due to batch, line and clone heterogeneity. Strategies to generate defined brain regions have advantages to disease modeling and screening, however the brain is ultimately multi-regional. Novel enhancements and engineering innovations are required to generate stereotyped multi-region brain organoids. Nonetheless, current protocols provide a new in-road to infer and perturb cell fate regulomes in human cerebral tissue.

## AUTHOR CONTRIBUTIONS

SJ, JSF generated organoids used in this study, with support from DW. SJ generated all single-cell transcriptome and accessible chromatin datasets with support from MSa. SJ performed the CROP-seq and the multiome experiments. JSF, DW, MSe constructed CROP-seq vectors. SJ, DW generated the isogenic GLI3 iPSC lines and generated the scRNA-seq data on GLI3 KO organoids. FZ, SJ performed western blots for GLI3. JSF performed the analysis of the scRNA-seq/scATAC-seq developmental time course with support from ZH, SJ. ZH performed receptor-ligand pairing analysis. JSF developed the Pando R package. JSF analyzed the CROP-seq data with support from ZH. SJ, JSF analyzed the GLI3 KO scRNA-seq and multiome data. JSF, SJ, ZH, JGC, BT designed the study and wrote the manuscript.

## DATA AVAILABILITY

Raw sequencing data will be deposited into ArrayExpress. Processed data is deposited on Zenodo (https://doi.org/10.5281/zenodo.5242913). The Pando R package is available on GitHub (https://github.com/quadbiolab/Pando). Other custom code used in the analyses is also deposited on GitHub (https://github.com/quadbiolab/organoid_regulomes).

## ACKNOWLEDGEMENTS

We thank the Camp, Treutlein, and Theis labs for helpful discussions. We thank Anne Weigert, Sabina Kanton, Theresa Schaffer and Ryoko Okamoto for assistance with stem cell and organoid culture and Tobias Gerber, Leila Sidow, and Keisuke Sekine for discussions relating to cloning. We thank Stephan Riesenberg and Svante Pääbo for kindly providing the iCRISPPR cell lines. We thank Michael Dannemann and Tomislav Maricic for providing genotype information for demultiplexing. Illumina sequencing was done by Barbara Schellbach and Antje Weihmann at the Max-Planck-Institute for Evolutionary Anthropology and Ina Nissen, Elodie Vogel Burcklen and Christian Beisel of the Genomics Facility at D-BSSE, ETH Zurich. FACS sorting support was provided by Mariangela Di Tacchio, Aleksandra Gumienny, Renan Antonialli and Thomas Horn of the single-cell facility at D-BSSE, ETH Zurich. We thank 10X Genomics for support with multiome experiments. No research with human ESC lines was funded by the ERC. This project has been made possible in part by, the Boehringer Ingelheim Fonds (J.S.F.), the Chan Zuckerberg Initiative DAF, an advised fund of Silicon Valley Community Foundation (CZF2019-002440, J.G.C, B.T.), European Research Council (Anthropoid-803441, J.G.C.; Organomics-758877, B.T.; Braintime-874606, B.T.), Swiss National Science Foundation (Project Grant-310030_184795, J.G.C; Project Grant-310030_192604, B.T.) and the National Center of Competence in Research Molecular Systems Engineering (B.T.).

## Methods

### Experimental methods

#### Stem cell and organoid culture

We used 3 human induced pluripotent stem cell (iPSC) lines (Hoik1, Wibj2 from the HipSci resource (37); 409B2 from the RIKEN BRC cell bank) and one human ES cell line (H9, WiCell). Stem cell lines were cultured in mTESR1 (Stem Cell Technologies, 05851) with mTeSR1 supplement (Stem Cell Technologies, 05852) and supplemented with penicillin/streptomycin (P/S, 1:200, Gibco, 15140122) on matrigel-coated plates (Corning, 354277). Cells were passaged 1-2 times per week after dissociation with TryplE (Gibco, 12605010) or EDTA in DPBS (final concentration 0.5mM) (Gibco, 12605010). Media was supplemented with Rho-associated protein kinase (ROCK) inhibitor Y-27632 (final concentration 5µM, STEMCELL, 72302) the first day after passage. Cells were tested for mycoplasma infection regularly using PCR validation (Venor GeM Classic, Minerva Biolabs) and found to be negative. 4,500 - 5000 cells were plated in ultra low attachments plates (Corning, CLS7007) to generate cerebral organoids using a whole brain organoid differentiation protocol (2). The use of human ESCs for the generation of cerebral organoids was approved by the ethics committee of northwest and central Switzerland (2019-01016) and the Swiss federal office of public health.

#### Single cell RNA and ATACseq for the developmental time course

Cerebral organoids were generated from four different stem cell lines (H9, 409B2, Wibj2, Hoik1) simultaneously. Cerebral organoids of the same batch were dissociated at multiple timepoints of the course of cerebral organoids development. We collected these single cell suspensions from an EB time point (day 4), the time points of neuronal induction (day 7, 9 and 11) and after embedding in matrigel and starting the neuronal differentiation process (day 12, 16, 18, 21, 26, 31 and 61). Organoids of the four different cell lines were pooled based on size and dissociated together and cell lines were later demultiplexed based on the single nucleotide polymorphism (SNP) information. Multiple organoids of each line were pooled together to obtain a sufficient number of cells. For the early time points 15 organoids per cell line were pooled, decreasing this number to minimally 3 organoids for the later time points (Data S1). Organoids were cut in halves and washed three times with HBSS without Ca2+/Mg2+ (STEMCELL technologies, 37250). Single cell suspensions were acquired by dissociation of the organoids with a papain-based dissociation (MACS Miltenyi Biotech 130-092-628). 2 ml of pre-warmed papain solution was added to the organoids and incubated for 15 minutes at 37 °C. Enzyme mix A was added before the tissue pieces were triturated 5-10 times with a 1000 wide-bore and p1000 pipette tips. The tissue pieces were incubated two times for 10 min at 37 °C with trituration steps in between and after with 200 and 1000p pipette tips. Cells were filtered with consecutively 30 µm and 20 µm pre-separation filters and centrifuged. Cells were resuspended and viability and cell count were assessed with a Trypan Blue assay on the automated cell counter Countess (Thermo Fisher). Cell suspensions were split in two and resuspended in CryoStor CS10 (STEMCELL technologies 07952) and cryopreserved at -80 °C. The next day cryotubes were transferred to liquid nitrogen for storage until the scRNA-seq and scATAC-seq experiments were performed. The cryopreserved single cell suspensions of each time point were thawed by warming up the cryo for 1-2 minutes in a water bath at 37°C and directly centrifuged in 10ml prewarmed DMEM with 10% FBS. Cells were washed twice with PBS + 0.04% BSA and filtered through a 40µm cell strainer (Flomi). For scATAC-seq, nuclei were isolated according to the protocol provided by 10x genomics (Demonstrated protocol CG000169 Rev D) using the low input protocol and a lysis time of 3 min. Nuclei were loaded in a concentration that would result in a recovery of 10,000 nuclei. In case of less nuclei recovered, the maximum number of nuclei was targeted. Single cell ATACseq libraries were generated with the Chromium Single Cell ATAC V1 Library & Gel Bead Kit. Prior to sequencing an additional clean-up step was performed to enrich shorter fragments by applying a double sided (1.2-0.75x) cleanup with AMPureXP beads (Beckman Coulter) and Illumina Free Adapter Blocking Reagent was used to reduce potential index hopping. The libraries were sequenced on Illumina’s NovaSeq platform. For sc-RNAseq, cells were put in a concentration after counting and viability checking that allowed targeting 10,000 cells and in case the cell number was not sufficient all cells were loaded. Single cell RNAseq libraries were generated with the Chromium Single Cell 3’ V3 Library & Gel Bead Kit. Single cell encapsulation and library preparation were performed according to the manufacturer’s protocol. Libraries were pooled, FAB treated and sequenced on Illumina’s NovaSeq platform. A summary of all single-cell experiments can be found in Data S1.

#### Doxycycline-inducible Cas9 nuclease and nickase cell line

The human iPSC line 409B2 was used to create an iCRISPR-Cas9 nickase (Cas9n) and an iCRISPR-Cas9 line as described (38). The doxycycline-inducible Cas9 expressing cell line was generated by introducing two transcription activator-like effector nucleases (TALENs) targeting the AAVS1 locus, which has shown to be effective for sustained transgene expression, and two TALEN constructs with donor plasmids. One of the donor plasmids contained a constitutive reverse tetracycline transactivator (AAVS1-Neo-M2rtTA) and the other one contained a doxycycline-inducible expression cassette (Puro-Cas9). A D10A mutation was introduced by site-directed mutagenesis of the original Puro-Cas9 donor with the Q5 mutagenesis kit (New England Biolabs, E0554S) to generate the Cas9n. The cell lines used were tested for the proper expression of pluripotency markers SOX2, OCT-4, TRA1-60, and SSEA, quantitative PCR confirmed the doxycycline-inducible Cas9n and digital PCR was used to exclude off-target integration (39). Both cell lines showed normal karyotypes upon generation, but the iCRISPR-Cas9 line acquired a common stem cell abnormality over time. 55% percent of the cells showed a derivative chromosome 2 with a long arm of chromosome 1 (bands q11q44) attached to the long arm of one chromosome 2 (band q37).

#### Vector and lentivirus preparation for perturbation experiment

The perturbation experiment was performed according to the CROP-seq protocol as described (17) with some minor alterations. The experiment was performed in organoids derived from the inducible Cas9 nuclease line which contains a Puro selection marker. To be able to select for cells that received the CROP-seq vector, Puro was exchanged for EGFP to isolate cells by fluorescence. Three gRNA per targeted gene were designed by Applied Biological Materials Inc. (abm) and synthesized by IDT as 74 base oligonucleotides with 19 and 35 bases of homology to the hU6 promoter and guide RNA backbone, respectively. Oligonucleotides were pooled in equal amounts and were assembled in the vector backbone by Gibson’s isothermal assembly. The plasmid library was sequenced to validate the complexity of the pooled plasmid library. 10ng of plasmid library was used for generating a sequencing library with a single PCR reaction. Illumina i7 and i5 indices were added by PCR and the library was sequenced on Illumina’s MiSeq platform. Upon validation, lentiviruses were generated by the Viral Core Facility of Charité Universitätsmedizin Berlin.

#### Generation of mosaic organoids for perturbation experiment

The iCRISPR-Cas9 line was cultured on matrigel in mTesr1 supplemented with P/S (1:200) and Cas9 was induced 2 days prior to lentiviral transduction by adding 2µg/ml doxycycline. 24 hours later cells were dissociated into single cells with TrypLE and 300.000 cells of the iCRISPR-Cas9 cells were plated in at least 12 wells of matrigel-coated 6-well plates in mTesr1 supplemented with P/S (1:200), Y-27632 (final concentration 5µM) and 2µg/ml doxycycline. 24 hours later cells were transduced with a low multiplicity of infection (MOI) where less than 30% of the cells were GFP+ to ensure that the majority GFP+ cells only received one lentivirus per cells. The viral particles were added to the culture media (mTesr1 supplemented with P/S, Y-27632 and 2µg/ml doxycycline). 24h later, media was exchanged for mTesr1 supplemented with P/S and 2µg/ml doxycycline until 70% confluency was reached. Cells were then sorted for GFP-positive to enrich for CROP-seq-vector-positive cells and plated on matrigel-coated plates in mTesr1supplemented with 100 µg/ml Primocin (InvivoGen, ant-pm-1) and Y-27632 (final concentration 5µM). When cells reached 70% confluency, whole brain organoids were generated as mentioned previously.

#### Preparation of single-cell transcriptomes from mosaic perturbed organoids

After 2 months, single organoids and pools of multiple organoids were dissociated with a papain-based dissociation kit (MACS Miltenyi Biotech, cat. No. 130-092-628) as described previously. Cells were sorted using fluorescence. Cell viability and number was assessed using the Trypan Blue assay and automated cell counter Countess (Thermo Fisher). Finally, cells were diluted to an appropriate concentration to obtain approximately 7,000 cells per lane of the 10x microfluidic chip. Single cell RNAseq libraries were generated with the Chromium Single Cell 3’ V3 Library & Gel Bead Kit. The expression libraries were FAB treated and sequenced on Illumina’s NovaSeq platform.

#### gRNA detection from single cell cDNA

gRNA were amplified from 60ng of cDNA remaining from scRNA-seq preparation with three separate PCR reactions similar to reactions as described (40). First, cDNA was amplified via PCR broadly targeting the outer part of the U6 promoter. Subsequently, the inner portion of the U6 promoter adjacent to the guide sequence and a TruSeq Illumina i5 adapter. Lastly, we added Illumina sequencing i7 adaptors. PCRs were monitored by qPCR to avoid over-ampliciation and following every PCR reaction the samples were purified using SPRI beads (Beckman Coulter) and libraries were sequenced at 1:10 proportion of the transcriptome library on Illumina’s NovaSeq.

#### gRNA detection from gDNA

Cells from different stages of the organoid protocol were collected (iPSC, embryoid body, embedded organoids and organoids day 30). QuickExtract 30-60 µl (Epicentre, QE0905T) was added to the cell pellets or organoids and the suspension was incubated at 65°C for 10 min, 68°C for 5 min and 98 °C for 5 minutes to extract DNA. The same PCR was done as used to validate the library complexity plasmid library (17). The PCR was performed with the KAPA2G Robust PCR Kit (Peqlab, 07-KK5532-03) using the supplied buffer B and 5 µl isolated DNA. The following program was used: 95°C 3 min; 35 ×(95°C 15s, 65 °C 15s, 72 °C 15s); 72°C 60s. Libraries were sequenced on with Illumina’s MiSeq (Nano kit).

#### Isogenic GLI3 knock-out line generation

Two days prior to lipofection, iPSC media was supplemented with 2µg/ml doxycycline (Clontech, 631311) to induce Cas9n expression. Two guides were designed using the Broad Institute’s CRISPR design tool (http://crispr.mit.edu/). The following guide pair was selected: ACAGAGGGCTCCGCCACGTGTGG, CCGCGGGACGGTGTTTGCCATGG. The Alt-R CRISPR-Cas9 System (IDT) was used for guide delivery with lipofection according to the manufacturer’s protocol. To form the crRNA:tracrRNA complex in a 3 µM final concentration for each guide complex, 1.5 µl of each guide crRNA was combined with 3 µl tracrRNA and 44 µl nuclease-free water. For the reverse transfection, 1.5µl of the crRNA:tracr complex mix and 0.75µl RNAiMAX (Invitrogen, 13778075) were diluted in 47.75 µl OPTI-MEM (Gibco, 1985-062) for each replicate and incubated for 20 minutes at RT in a well of 96-well plate coated with Matrigel (Corning, 35248). During incubation, ∼70% confluent cells were detached with TryplE (Gibco, 12605010), centrifuged and resuspended in 1ml mTeSR with 1:1000 Y-27632 (STEMCELL, 72302). After complex incubation, cells were diluted 30 or 60 times in 100 µl mTeSR with 1:1000 Y-27632 (STEMCELL, 72302) and 2µg/ml doxycycline (Clontech, 631311) and the cell suspension was added to a well containing the transfection complexes. After 24h media was replaced with mTeSR1 media and cells were allowed to recover for 72h. 70% confluent wells were used for further processing after 72 hours. Cells were passaged as single cells in a Matrigel-coated (Corning, 35248) 6-well plate in mTeSR media with 1:200 P/S (Gibco, 15140122) and 1:1000 Y-27632 (STEMCELL, 72302). Low amounts of cells were plated per well to avoid the fusion of colonies. Media was changed daily and Y-27632 was added for the first 72 hours to prevent apoptosis of the single cells. When colonies were apparent, single colonies were picked by scraping with a 10 microliter pipette tip. 2/3 of the cell suspension was plated in a single well of a Matrigel-coated 96-well plate in mTeSR1 supplemented with 1:200 P/S and 1:1000 Y-27632. The other portion of the cell suspension was pelleted and used for validation of frameshift mutations by sequencing. Validated clones were expanded, cryopreserved and karyotyped. The three selected lines, one WT and 2 KO lines, showed a normal karyotype.

#### Validation of KO lines by sequencing

The cell pellets of picked colonies were resuspended in 10 µl QuickExtract (Epicentre, QE0905T) and the suspension was incubated at 65 °C for 10 min, 68° C for 5 min and 98 °C for 5 minutes to extract DNA. A PCR was performed with primers containing Illumina sequencing adapters for the targeted locus of the GLI3 gene. Amplification was performed with the KAPA2G Robust PCR Kit (Peqlab, 07-KK5532-03) using the supplied buffer B and 2 µl of extracted DNA. The following program was used: 95 °C 3 min; 35 ×(95°C 15s, 65 °C 15s, 72 °C 15s); 72°C 60s. Unique P5 and P7 Illumina indices were added to 0.5 µl of the previous PCR product with a second PCR program (98°C 30s; 25 ×(98°C 10s, 58°C 10s, 72°C 20s); 72°C 5 min), with the Phusion HF MasterMix (Thermo Scientific, F-531L). The double-indexed libraries were pooled and purified with SPRI beads. Purified libraries were sequenced on the MiSeq (Illumina) resulting in pair-end sequences of 2 × 150 bp. LeeHom (41) was used to trim the adapters after base calling with Bustard (Illumina).

#### Western blot

GLI3 WT and KO organoids of day 15 were collected into Laemmli-Buffer, homogenized with a pestle (Fisher-brand 12-141-368) and sonicated with 15 cycles with the Bioruptor Plus. Subsequently, two high and low amounts of protein extractions and ladder (Thermo Scientific 26620) were run on 8% SDS-PAGE (BioRad System) and transferred to PVDF membrane using Wet-Blot. After blocking for 20 min with 4% milk powder in PBS+0.1% Tween, the primary antibody (1:1000, stock 0.5 µg/µl, R&D systems, AF3690) was incubated overnight at 4°C. After washing 3 times for 7 min at RT in PBS+0.1% Tween on a shaker, the secondary Goat IgG HRP-conjugated antibody (1:7000, R&D systems HAF017) diluted 1:5000 in 4% milk and PBS+0.1% was incubated for 2h. The enhanced chemiluminescence signal was recorded using ChemiDoc. The loading control Beta-Catenin (Primary antibody: stock 1:10.000, Cell Signalling technologies L54E2; Secondary antibody: stock 0.8 µg/µl 1:7000, Jackson ImmunoResearch 115-035-003) was probed on the same membrane and loading was also controlled by ponceau staining. Raw images are provided in Data S8.

#### Generation of single cell transcriptome and multiome of GLI3 KO and WT organoids

Organoids of GLI3 WT and KO iPSCs were generated simultaneously and dissociated with a papain-based dissociation kit (MACS Miltenyi Biotech, cat. No. 130-092-628) as described earlier. ScRNAseq was performed on day 45 of organoid development for both KO lines and the WT line for two independent organoid batches. After dissociation, cell viability was checked, cells were counted and 7000 cells were targeted per lane of the 10x microfluidic chip. Libraries were generated with the Chromium Single Cell 3’ V2 Library & Gel Bead Kit and sequenced on Illumina’s HiSeq platform. Combined scRNA-seq and scATAC-seq were generated with the Chromium Single Cell Multiome ATAC + Gene Expression kit. Prior to nuclei isolation, day 19 organoids were dissociated with the papain-based dissociation. Nuclei were isolated according to the protocol provided by 10x genomics (Demonstrated protocol CG000365, Rev B) with a lysis time of 3 min. The gene expression and accessibility libraries were FAB treated and sequenced on Illumina’s NovaSeq platform.

### Data analysis methods

#### Preprocessing of single-cell RNA-sequencing data from the organoid time course

We used Cell Ranger (version 3.0.2) with default parameters to obtain transcript count matrices by aligning the sequencing reads to the human genome and transcriptome (hg38, provided by 10x Genomics, version 3.0.0). Count matrices were further preprocessed using the Seurat R package (version 3.2) (19). First, cells were filtered based on unique molecular identifier (UMI) counts, the number of detected genes, and the fraction of mitochondrial genes. The threshold of mitochondrial gene fraction was held constant across datasets (<0.2). Since sequencing depth varied between timepoints, the threshold of UMI count and number of detected genes was set individually for each sample as follows:

- Day 4&7: #UMI: >10000, <80000; #features: >3000, <8000
- Day 11: #UMI: >10000, <60000; #features: >3000, <8000
- Day 12: #UMI: >2500, <40000; #features: >1000, <6000
- Day 16: #UMI: >10000, <60000; #features: >3000, <8000
- Day 18&21: #UMI: >2500, <60000; #features: >1500, <8000
- Day 26: #UMI: >2500, <60000; #features: >2000, <8000
- Day 31: #UMI: >2500, <50000; #features: >2400, <7500
- Day 61: #UMI: >1000, <60000; #features: >1000, <8000

Transcript counts were normalized by the total number of counts for that cell, multiplied by a scaling factor of 10000 and subsequently natural-log transformed (`NormalizeData()`).

#### Preprocessing of single-cell ATAC-sequencing data from the organoid time course

We used Cell Ranger ATAC (version 1.1.0) with default parameters to obtain fragment files by aligning the sequencing reads to the human genome and transcriptome (hg38, provided by 10x Genomics, version 1.1.0). Peaks were called from the fragment file using MACS2 (version 2.2.6). Both the fragment files and the peak count matrices were further preprocessed using Seurat (version 3.2) (19) and Signac (version 1.1) (42). First, peaks were filtered by width (<10000 bp, >20 bp) to retain only high-quality peaks. Further, the following quality control metrics were computed using Signac: A transcription start site (TSS) enrichment score (`TSSEnrichment()`), nucleosome signal (`NucleosomeSignal()`), the percentage of reads in peaks, and the ratio of reads in genomic blacklist regions. Subsequently, cells were filtered based on the following metrics:

- Percentage of reads in peaks > 30%
- # peak region fragments > 5000
- Blacklist ratio < 0.003
- Nucleosome signal < 5
- # TSS fragments > 5000
- TSS enrichment score > 2

We then created a unified set of peaks from the union of peaks from all samples by merging overlapping and adjacent peaks. The unified set of peaks was requantified for each sample using the fragment file (`FeatureMatrix()`). Peak counts were normalized by term frequency–inverse document frequency (tf-idf) normalization using the Signac functions `RunTFIDF()`.

#### Demultiplexing of different lines based on SNV information

Cells pooled from different stem cell lines were demultiplexed using demuxlet (43). Genotyping information was called using bcftools based on DNA-seq data (H9 and 409B2) (44) or downloaded from the HipSci website. All files were merged using bcftools and sites with the same genotypes in all samples were filtered out. Demuxlet was run with default settings. Cells with ambiguous or doublet assignments were removed from the data. For all other cells the best singlet assignment was considered the line label.

#### Integration of transcriptome and chromatin accessibility data

To create a shared feature space between the two modalities, gene activities were calculated from chromatin accessibility data using the Signac function `GeneActivity()` with default parameters and subsequently log-normalized. For each time point and line separately, we performed Canonical Correlation Analysis (CCA) on gene activities and gene expression data using the Seurat function `RunCCA()` based on 2000 features, which were selected using the Seurat function `SelectIntegrationFeatures()`. In CCA space, we performed minimum-cost maximum-flow (MCMF) bipartite matching between the modalities as described (20) (https://github.com/ratschlab/scim). The function `get_cost_knn_graph()` was used with `knn_k=10`, `null_cost_percentile=99`, `capacity_method=‘uniform’` and otherwise default parameters. Based on the bipartite matches, matched cells were summarized to metacells containing measurements from both modalities. If multiple cells from one modality were included in a metacell, the arithmetic mean between cells was calculated.

#### Removal of cells with glycolysis signature

An additional quality control step was applied on the level of metacells to remove cells with transcriptomic signatures of glycolysis upregulation. This was based on primary cell-type predictions by using public human fetal brain scRNA-seq data (Nowakowski dataset) (45). We fit a multinomial logistic regression model with lasso regularization penalty (`alpha=1`), using gene-expression ranks of the Nowakowski dataset as the training set, to predict the cell-type identity of metacells in the organoid developmental time course. Metacells which were predicted to be of ‘glycolysis’ identity were excluded from the dataset. To fit the logistic regression model and automatically determine the regularization parameter lambda through cross-validation, we used the function `cv.glmnet()` from the glmnet R package.

#### Integration of different lines and timepoints

Integration of lines and timepoints was performed using the log-normalized gene expression data of metacells. To select a set of features suitable for integration of all lines and timepoints, we selected the union of the 100 most variable genes for each timepoint separately (local) as well as across the full dataset (global). Analogously, we selected the union of locally and globally variable transcription factors (Data S2). We used the union of the selected genes and TFs and further excluded cell cycle related genes (46) from the set. Next, we computed cell cycle scores using the Seurat function `CellCycleScoring()`. Subsequently the data was z-scaled, cell cycle scores were regressed out (`ScaleData()`) and Principal Component Analysis (PCA) was performed using the Seurat function `RunPCA()`. We used the first 10 principal components (PCs) to integrate the different timepoints in the dataset using the Cluster Similarity Spectrum method (CSS) (21). To remove any remaining signal cell cycle signal for any downstream tasks, we again regressed out the cell cycle scores from the integrated CSS matrix. To obtain a two-dimensional representation of the data we performed Uniform Manifold Approximation and Projection (UMAP) (47) using `RunUMAP()` with `spread=0.5`, `min.dist=0.2` and otherwise default parameters.

#### Calculation of motif enrichment scores

Position weight matrices (PWM) of human TF binding motifs were obtained from the CORE collection of JASPAR2020 (48). Motif positions in accessible chromatin regions were determined using the R package motifmatchr (version 1.14) (https://doi.org/10.18129/B9.bioc.motifmatchr) through the Signac function `FindMotifs()`. Enrichment scores of motifs in accessible regions were calculated for each metacell using chrom-VAR (49) through the Signac function `RunChromVAR()`.

#### RNA velocity calculation

To obtain count matrices for the spliced and unspliced transcriptome, we used kallisto (version 0.46.0) (50) by running the command line tool loompy fromfastq from the python package loompy (version 3.0.6)(https://linnarssonlab.org/loompy/). Spliced and unspliced transcriptomes were summarized to metacell level as described above. RNA velocity was subsequently calculated using scVelo (version 0.2.2) (27) and further analyzed using scanpy (version 1.7.0) (51). First, 2000 highly variable features were selected using the function `scanpy.pp.highly_variable_genes()`. Cell cycle genes (46) were excluded from this feature set and the dataset was subsetted to the resulting gene set. Subsequently, moments were computed in CSS space using the function `scvelo.pp.moments()` with `n_neighbors=30`. RNA velocity was calculated using the function `scvelo.tl.velocity()` with `mode=‘stochastic’` and a velocity graph was constructed using `scvelo.tl.velocity_graph()` with default parameters. To order cells in the developmental trajectory, a root cell was chosen randomly from cells of the first time point (day 4) and velocity pseudotime was computed with `scvelo.tl.velocity_pseudotime()`. The obtained velocity pseudotime was further rank-transformed and divided by the total number of metacells in the dataset.

#### Annotation of organoid developmental stages

To annotate different organoid developmental stages, we first divided the dataset in 20 bins based on quantiles of velocity pseudotime. For each bin, we computed the average gene expression and peak accessibility across metacells and computed the pairwise Pearson correlation between log-normal gene expression values of each bin. From the correlation coefficient r, we defined a distance metric as 1-rand used it to perform hierarchical clustering using the `ward.D2` method as implemented in the stats R package (`hclust()`). Based on the resulting clusters, bins were manually annotated as pluripotent (PSC), neuroectoderm, neuroepithelium, neural progenitor (NPC) or neuron.

#### Identification of stage-specific chromatin access

To find sets of peaks with stage-specific accessibility, we computed for each stage the percentage of metacells in which each peak was detected. We then computed a specificity score through dividing the detection percentage for each stage by the detection percentage of all other metacells. We filtered peaks with in-stage detection percentage of > 15% and a stage specificity of >1.5. From these peaks we selected the top 5000 peaks with the highest specificity score. Using these specific peak sets for each stage, we used GREAT (35) with the GRCh38 genome assembly and otherwise default parameters to obtain functional enrichment results. We reported GO Biological Process enrichments with FDR < 0.01 and that were supported by >30 foreground regions.

#### Correlation with organizer populations in the developing mouse brain

To assess if certain populations in the organoid developmental time course resembled potential organizer populations, we compared the cell populations arising in the data with organizer populations from a developmental mouse brain atlas (24). The gene expression data data was obtained from http://mousebrain.org/ and subset to annotated organizer populations. To perform comparisons to human transcriptomics data, we converted gene names (official gene symbols) to the names of the human homolog, using homology annotations from the Mouse Genome Informatics database (http://informatics.jax.org). We selected organizer-specific genes by performing differential expression between organizer populations (Wilcoxon rank-sum test, FDR < 0.01) and selecting the top 30 genes with the highest fold change for each organizer (organizer markers, Data S2). We then subset our dataset into different, partially overlapping stages, representing interesting transition points (neuroectoderm emergence: day 7-11; neuroepithelium emergence: day 12-21; branching stage: > day 11, between 15% and 85% pseudotime quantile). To better resolve potential organizer populations each subset was re-integrated by performing CSS as described above based on the union of variable features (100 for neuroectoderm and neuroepithelium, 400 for branching stage, Data S2) and organizer markers. Each subset was clustered using the Louvain algorithm (52) in CSS space with a resolution of 0.8 using the Seurat function `FindClusters()`. We calculated the average log-normal expression values for each cluster and mouse organizer and then computed the Peason correlation between cluster and organizer transcriptomes based on organizer markers.

#### Cell-cell communication analysis

To investigate ligand-receptor mediated cell-cell communications in developing cerebral organoids, we focused on the branching stage subset of neural progenitor cells as described above. CellPhoneDB (version 2.0) (53) with default setting was applied to the single-cell transcriptomic profiles and the Louvain clustering results described above (14 clusters in total). This analysis identified ligand-receptor (LR) pairs which significantly co-expressed in pairs of cell clusters, and therefore potentially mediated the communications between cell populations (permutation test, P<0.05). For each significant LR pair between one cell cluster pair, the cell cluster expressing the ligand gene was considered as the signal source, and the one expressing the receptor gene was considered as the signal target. To identify groups of ligand-receptor pairs mediating similar cell-cell communications, a hierarchical clustering was applied to the identified significant LR pairs using COMUNET (54), based on their signaling patterns in different cell cluster pairs.

#### Inference of regional cell-fate trajectories

To resolve the regional cell fate branches we relied on CellRank (version 1.3.0) to compute transition probabilities into terminal cell states and PAGA (55) to obtain a graph abstraction of the transcriptomic manifold. First, terminal neuronal states were annotated manually using VoxHunt (version 1.0.0) (5) based on the top 20 structure markers. To resolve the developmental trajectories leading up to the emergence of neurons with distinct regional identities, transition probabilities to each of the terminal states were computed for each cell using CellRank. A transition matrix was constructed by combining a velocity kernel (`VelocityKernel()`) and a connectivity kernel (`ConnectivityKernel()`) with weights of 0.5 each. Absorption probabilities for each of the predefined terminal states were computed using the GPCCA estimator. From these probabilities, we computed a transition score by ranking the absorption probabilities and normalizing by dividing by the total number of metacells. We then constructed a graph abstraction of the dataset by high-resolution clustering using the Louvain algorithm (52) with a resolution of 20. We used PAGA to compute the connectivities between clusters (`scvelo.tl.paga()`) and summarized transition scores for each of the clusters. To find branch points at which the transition probabilities into different fates diverge, we then constructed a nearest-neighbor graph between the high-resolution clusters based on their transition scores (`k=30`). We further pruned the graph to only retain edges between nodes with a connectivity score of > 0.2 and edges going forward in pseudotime, i.e. from a node with a lower velocity pseudotime to a node with a higher velocity pseudotime. The resulting graph is directed with respect to pseudotemporal progression and represents a coarse-grained abstraction of the fate trajectory, connecting groups of cells with both similar transition probabilities to the different lineages and high connectivities on the transcriptomic manifold. To assign fate identities to each branch in the graph, we first selected the nodes with the highest transition probability and pseudotime for each of the terminal states as tips. We then performed 10000 random walks with 200 steps from each tip along edges backwards in pseudotime using the igraph R package (version 1.2.6) (https://igraph.org/). Next, we computed for each node the visitation frequency from each of the terminal states. We then assigned branch identities to each node based on the visitation frequencies as follows: If a node’s visitation frequency from one tip was more than 100x higher than from the next highest tip, it was unambiguously assigned the identity of this tip. If the visitation frequencies from multiple tips were within 100x of each other, then the node was assigned the identity of all of such tips. Nodes that were assigned both the dorsal telencephalic and ventral telencephalic identity were relabelled as ‘telencephalon’. Nodes that were assigned all three identities were labelled as ‘early’ to indicate that their fate was not yet committed. Nodes that could not be reached through this procedure were assigned the identity of the node with the highest connectivity score. The final labelled graph was visualized using the Fruchterman-Reingold layout algorithm as implemented in the igraph R package.

#### Gene regulatory network inference with Pando

We developed Pando to infer gene regulatory networks while taking advantage of multi-modal single-cell measurements, where both the RNA and the ATAC components are either measured for each cell or integrated to obtain metacells or clusters with both modalities. The core GRN inference algorithm in Pando can be summarized in four main steps:

1. Selecting candidate regulatory genomic regions
2. Scanning regions for transcription factor binding motifs
3. Selecting region-TF pairs for each target gene
4. Constructing a linear model with region-TF pairs as independent variables and the expression of the target gene as the response variable The coefficients of this linear model can now be seen as a measure of interaction between the region-TF pair and the downstream gene, resulting in a regulatory graph. In the following, we will describe these steps in more detail.

#### Selection of candidate regions for GRN inference

To narrow the set of genomic regions that are taken into account for each target gene when constructing the model, we can take advantage of prior knowledge about the potential importance of these regions. Genomic sequence conservation is one such criterion that indicates the relevance of a stretch of DNA, as it has been maintained by natural selection. Thus, we first intersected the peak regions in the ATAC-seq data with the set of PhastCons conserved elements (56) from an alignment of 30 mammals (obtained from https://genome.ucsc.edu/). As exonic regions tend to be conserved regardless of their regulatory relevance, we further excluded exonic regions from this set. Further, we considered candidate cis-regulatory elements (cCREs) derived from the ENCODE project (29). For this, we obtained the set of all human cCREs from https://screen.encodeproject.org/ (GRCh38) and intersected it with peak regions. The union of the resulting conserved and cCRE regions was carried forward as the set of candidate regions for GRN inference.

#### Construction of an extended motif database for GRN inference

Because TFs need to be matched with potential binding sites, the availability of a binding motif is required for a TF to be included in the GRN. Therefore, we aimed to gather motif information for all TFs relevant in our dataset. First, we selected the union of the 4000 most variable genes in each individual time point (Data S2). All TFs in this set were considered relevant. We then obtained binding motifs from JASPAR (2020 release) (48) taking into account the CORE and the UNVALIDATED collection. For TFs where no binding motif was available in JASPAR, we further considered the CIS-BP database (57). Where possible, motifs with direct experimental evidence were prioritized over inferred motifs and motifs that were inferred based on other JASPAR motifs were prioritized over the rest. For all relevant TFs that were also not covered by CIS-BP, motifs were inferred based on protein sequence similarity to other TFs from the same family. Family information and protein sequences for all TFs were obtained from AnimalTFDB (58) and pairwise multiple sequence alignments were performed using the Needleman-Wunsch algorithm (59) as implemented in needle from the EMBOSS software suite (version 6.5.7) (60). For each query TF, we considered all TFs from the same family with a global sequence similarity of at least 20% and selected the motifs from the 3 most similar TFs. TF motifs from all sources were combined into one database and motif positions in accessible chromatin regions were determined using the R package motifmatchr (version 1.14) (https://doi.org/10.18129/B9.bioc.motifmatchr) through the Signac function `FindMotifs()`.

#### Coarse-graining expression and chromatin accessibility data

Before inferring the GRN, we coarse-grained the data to denoise and remove sparsity. First, we summarized the expression and chromatin accessibility of close cells using the pseudocell algorithm outlined in (44). In brief, we randomly selected 30% of all cells in the dataset as the seed cells and constructed a territory for each seed with the 10 nearest neighbors based on euclidean distances using the top 20 PCs. If one cell was assigned to multiple territories, one was randomly chosen. For all cells contained in a territory, gene expression data was summarized using the arithmetic mean. For chromatin accessibility data, an accessibility probability for each territory was computed by averaging binarized read counts. We further performed Latent Semantic Indexing (LSI) on the peak counts of each territory using the Signac functions `RunTFIDF()` followed by `RunSVD()`. Based on the top 20 LSI components we further performed high-resolution clustering using the Louvain algorithm with a resolution of 100 and accessibility probabilities were further summarized to a cluster level by computing the arithmetic mean so that each cell in the cluster was represented by the same vector.

#### Linear model-based GRN inference

Pando used a linear model-based approach to infer the regulatory interactions between TF-binding site pairs and the corresponding gene. Genomic coordinates for all genes were obtained via the R package EnsDb.Hsapiens.v86 (https://doi.org/10.18129/B9.bioc.EnsDb.Hsapiens.v86). For each gene, we considered a regulatory region encompassing the gene body and 100 kb upstream of the transcription start site (TSS). We then define a linear model on the (log-normalized) expression Y of the gene i based on all TF-binding site interactions in this region:

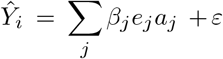

Where *e*_*j*_ is the log-normalized expression of transcription factor *j, a*_*j*_ is the accessibility probability of the peak that overlaps its binding region, *β*_*j*_ is the fitted coefficient for this interaction and *ε* is the intercept. The fitted coefficients can then be interpreted as the regulatory effect of TF-binding site pairs on the downstream genes. To fit the linear model, we use the function `glm()` from the stats R package using gaussian noise and an identity link function.

#### Peak and gene module construction

To prune the network and retain only significant interactions, the fitted coefficients were tested for statistical significance using ANOVA. We corrected for multiple testing using the Benjamini-Hochberg method to obtain an FDR-adjusted p value, to which a significance threshold of 0.05 was applied. The remaining connections were further summarized to extract sets of negatively (coefficient < 0) and positively (coefficient > 0) regulated target genes and regulatory regions for each transcription factor.

#### Pando implementation details

Pando was implemented as an R package and is available on GitHub (https://github.com/quadbiolab/Pando). Pando was designed for easy use and integrates smoothly with widely used single-cell analysis tools in R, namely Seurat and Signac. Its core functionality is implemented in four main functions:

`initiate_grn()` selects candidate regions from the dataset and initiates the object for GRN inference. The user can flexibly define custom sets of candidate regions to be taken into account by Pando.

`find_motifs()` scans candidate regions for transcription factor motifs. The motif database constructed in this work is included in the pando package, but can also be manually supplied.

`infer_grn()` selects regulatory regions for each target gene and performs the linear model fitting. We implemented support for all generalized linear models provided by the stats R package as well as regularized linear models provided by the glmnet R package (version 4.0).

`find_modules()` constructs gene and regulatory modules for each transcription factor.

The implementation is flexible and allows the user to apply the Pando framework to a wide range of use-cases.

#### Visualization of the GRN

We sought to visualize the inferred transcription factor network based on both co-expression and regulatory relationships between transcription factors. First, we computed the Pearson correlation between log-normalized expression of all transcription factors in the network across all metacells in the atlas. From the correlation value r and estimated model coefficient ß between all transcription factors i and j, we then computed a combined score s as

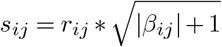

resulting in a TF by TF matrix. We performed PCA on this matrix and used top 20 PCs as an input for UMAP as implemented in the uwot R package (https://github.com/jlmelville/uwot) with default parameters.

#### Selecting transcription factors with region-specific expression

We used a discriminative learning approach to select transcription factors whose expression was specific to a regional branch. Using the dataset with 2 month old organoids (44), we fit a multinomial logistic regression model with elastic net regularization (alpha=0.5) to distinguish the different regional branches based on log-normalized gene expression. Genes with non-zero coefficients for a given branch were considered region-specific.

#### Calculation of module activity and analysis of module branch specificity

Based on the GRN inferred by Pando, the activity of a transcription factor can be represented by the expression of the set of genes it regulates (gene modules) or by the accessibility of its set of regulatory regions (regulatory modules). To calculate the activity of gene modules we used the Seurat function `AddModuleScore()` with all genes included in GRN inference as the background (pool). For regulatory modules, we used the R package chromVAR (version 1.14) (49) to obtain a set of background peaks (`getBackgroundPeaks()`). We then computed deviations in accessibility from the background for each regulatory module (`computeDeviations()`). Next, we assessed how the activity of positively regulated gene modules varied during neurogenesis over pseudotime and between branches. For this analysis we excluded all cells from the PSC and neuroectoderm stage. We fit three gaussian linear models for each gene i module with module activity (Y) as the response variable and branch assignment and/or velocity pseudotime as the independent variables:

1. *Y*_*i*_ ∼ *branch*
2. *Y*_*i*_ ∼ *pseudotime*
3. *Y*_*i*_ ∼ *branch* + *pseudotime*

We used the *R*^2^ value of these models as the fraction of variance explained by branch (1), pseudotime (2) or branch and pseudotime (3). We further tested for differential module activity between the branches for each branch point separately using a Wilcoxon rank-sum test as implemented in the R package presto (61). For the comparison of dorsal and ventral telencephalon, we only considered cells in the top 30% pseudotime quantile (NPC and Neuron stages). To visualize dorsal and ventral telencephalon-specific transcription factor networks, we first selected positively regulated gene modules of transcription factors with branch-specific expression (described above). For each branch, we then selected the top 15 modules whose module activity was significantly upregulated (FDR<0.05) based on the mean difference of module activity between the branches.

#### Preprocessing, integration and annotation of CROP-seq single-cell RNA-sequencing data

As with the organoid time course, count matrices were obtained with Cell Ranger (version 3.0.2) and further preprocessed using the Seurat R package (version 3.2) (19). First, cells were filtered based on unique molecular identifier (UMI) counts (>500, <30000), the number of detected genes (>500, <6000) and the fraction of mitochondrial genes (<0.1). Transcript counts were normalized by the total number of counts for that cell, multiplied by a scaling factor of 10000 and subsequently natural-log transformed (`NormalizeData()`). The different samples were integrated using RSS (44) based on the 2000 most variable features (`FindVariableFeatures()`). In RSS space, we performed Louvain clustering with a resolution of 3. Regional identities as well as NPC/Neuron identities were assigned to Louvain clusters using a combination of VoxHunt similarity maps and canonical marker genes. Cells annotated as off-target cell types such as mesenchyme and choroid plexus were removed from all downstream analyses.

#### Assignment of gRNA labels to cells

To assign gRNA labels to cells, reads obtained from amplicon sequencing were first aligned to the human genome and transcriptome (hg38, provided by 10x Genomics), which was extended with artificial chromosomes representing the CROP-seq-Guide-GFP construct (17), using Cell Ranger. We observed that read counts of gRNA UMIs followed a bimodal distribution, with the lower peak likely representing sequencing or amplification artifacts. To extract the higher peak, we first fit a Gaussian Mixture Model with two components on natural log-transformed read counts using the function `GaussianMixture()` from the scikit-learn python package (https://scikit-learn.org/). We then used a probability cutoff of 0.5 to extract the mixture component with higher average read counts. From these gRNA UMIs we constructed a cell x guide count matrix, which was further binarized to obtain the final cell-to-gRNA assignments.

#### Inference of perturbation probability

To account for a potential mixture of unperturbed and perturbed cells in the population, we inferred probabilities of a gRNA having a phenotypic effect on the cell using the strategy previously proposed (15). Here, a Bayesian approach is used to obtain the probability of a cell being perturbed given the observed transcriptome. To this end, a regression model is fit with the gene expression matrix as the response Y and the native gRNA assignments, cell and sample covariates as independent variables (X):

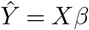

After fitting the model, the model fit is re-evaluated for each cell with the gRNA assignment set to 0 (*X*_0_).The difference of the squared errors of the two fits can then be transformed into a probability with:

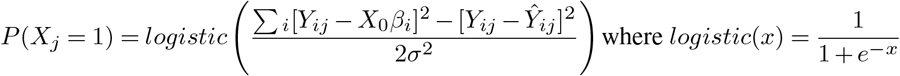

As in the original publication, we used linear regression model with elastic net regularization (`alpha = 0.5`) using gaussian noise and an identity link function to fit the model on Y=X based on 500 most variable features. The regularization parameter lambda was automatically determined through cross-validation as implemented in the function cv.glmnet() from the glmnet R package. Models were fit for each gene i on log-normalized transcript counts Y with binary assignments X for each gRNA j as well as celltype, sample and number of detected genes as covariates:

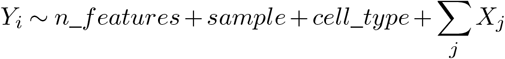

After computing the above described perturbation probabilities for each cell and gRNA, they were further summarized to a target gene level by taking the maximum probability among the three gRNAs targeting the same gene.

#### Determination of transcriptomic knock-out (KO) effects in the CROP-seq screen

To determine how gene KOs affect the transcriptomic state of neuronal populations arising in brain organoids we used a linear model-based approach (15). For each neuronal type, we inferred perturbation probabilities p for each target gene j as mentioned above and fit a linear model on log-normal transcript counts (Y) for each gene i as follows:

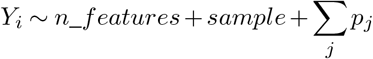

To determine KO effects in neural NPCs, we additionally used cell cycle phase as a covariate. For this we inferred the cell cycle phase with the Seurat function `CellCycleScoring()` and then constructed the linear model as follows:

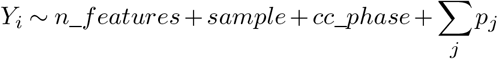

To determine the KO effects across all neurons, we inferred global perturbation probabilities on the full dataset and then fit a linear model across neuronal populations on log-normal transcript counts for each gene y as follows:

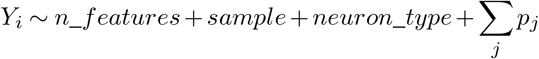

The coefficients for each target gene were tested using ANOVA and multiple-testing correction was performed using the Benjamini-Hochberg method to obtain a FDR-adjusted p value. Genes for which the coefficient of a target gene were significant (FDR < 10^−4^) were treated as differentially expressed genes for this target gene.

#### Determination of composition changes in the CROP-seq screen

To assess the degree to which the knock-out of a target gene changes the regional composition of the organoid, we first tested the enrichment of each gRNA in each regional branch. To control for confounding effects through differential gRNA abundance in different organoids, we used a Cochran-Mantel-Haenzel (CMH) test stratified by organoid. Additionally, we performed a Fisher’s exact test to test for enrichment for each organoid individually. Multiple-testing correction was performed using the Benjamini-Hochberg method. To account for other potential within-sample confounders such as clonal heritage, we first required for each gRNA that the enrichment was significant (FDR < 0.05) in more than one individual organoid and that the direction of each significant enrichment was consistent across organoids. All gRNAs for which this was not the case were removed. In a second step, we further required for remaining gRNAs that the same significant effect (FDR < 0.01) was observed for at least one other gRNAs targeting the same gene. For the remaining gRNAs we summarized the assignments for each target gene *i* and calculated the log odds ratio of the enrichment in each regional branch j with

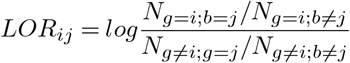

where N a matrix of cell counts for each target gene in each branch. For each target gene, the maximum log odds ratio across the three branches was treated as a measure of composition change.

#### Preprocessing and integration of single-cell

RNA-sequencing data from the GLI3 knock-out experiment Transcript count matrices were obtained with Cell Ranger (version 3.0.2) and further preprocessed using the Seurat R package (version 3.2) (19). First, cells were filtered based on unique molecular identifier (UMI) counts (>200, <60000), the number of detected genes (>200, <6000) and the fraction of mitochondrial genes (<0.1). Transcript counts were normalized by the total number of counts for that cell, multiplied by a scaling factor of 10000 and subsequently natural-log transformed (`NormalizeData()`). From all protein coding, non-mitochondrial and non-ribosomal genes we selected the 200 most variable based on the vst method (`FindVariableFeatures()`). PCA was performed based on the z-scaled expression of these features. Different samples were integrated using CSS (21) based on the top 20 PCs with default parameters. To visualize the dataset in two dimensions, we used UMAP on the CSS coordinates with `spread=0.5, min.dist=0.2`.

#### CRISPResso analysis and protein sequence prediction

To find clones with a frame-shift mutation, CRISPResso was used to analyse the sequencing data (62). This tool aligned the amplicons to the wild-type gene sequence to call inframe and frameshift indels. Analyses were performed with the following parameters: `-w20`, `-min_indentity_score70`, and `-ignore_substitutions`. Substitutions were ignored, only sequences with a minimum of 70% similarity were used and only indels present in a window of 20 base pairs from each of the gRNAs were called. Cell lines were considered as KO when > 98% of the reads were considered as a non-homologous end-joining event, the indels caused a frameshift, not more than two different indels were seen and were present in a 50:50 distribution. The predicted protein sequence was obtained with the Biopython python package (63).

#### Preprocessing and integration of multiome data from the GLI3 knock-out experiment

Initial transcript count and peak accessibility matrices were obtained with Cell Ranger Arc (version 1.0.0) and further preprocessed using the Seurat (version 3.2) (19) and Signac (version 1.1) (42) R packages. Peaks were called from the fragment file using MACS2 (version 2.2.6) and combined in a common peak set before merging. Cells were filtered based on transcript (UMI) counts (>1000, <25000), mitochondrial transcript percentage (<30%), peak fragment counts (>5000, <700000) and TSS enrichment score (>1). Transcript counts were normalized by the total number of counts for that cell, multiplied by a scaling factor of 10000 and subsequently natural-log transformed (`NormalizeData()`). Principal Component Analysis (PCA) was performed using the Seurat function `RunPCA()`. Different samples were integrated based on the top 20 PCs with Harmony (64) using the function `RunHarmony()` from the R package SeuratWrappers (version 0.3.0)(https://github.com/satijalab/seurat-wrappers) with max.iter.harmony=50 and otherwise default parameters.

#### Annotation of cells from the GLI3 knock-out experiment

To annotate the cell states from both the scRNA and the multiome experiments, we made use of the annotations of the annotated multi-omic atlas of organoid development that was previously generated. We transferred the regional branch labels using the method implemented in Seurat using the functions `FindTransferAnchors()` and `TransferData()`. We then performed Louvain clustering with a resolution 1 for the scRNA data and 0.8 for the multiome data. Clusters were manually assigned to branch identities based on the transferred labels as well as marker gene expression. In case of the multiome data, we identified populations of mesenchymal and non-neural ectoderm cells, which were excluded from the downstream analysis.

#### Differential expression analysis for the GLI3 knock-out experiment

To assess the transcriptomic effects of the GLI3 knock-out in ventral telencephalon neurons, we performed differential expression analysis using a linear model-based approach analogous to the approach used in the CROP-seq screen. We fit a linear model on log-normal transcript counts *Y* for each gene *i* with the knock-out label and number of detected features as independent variables:

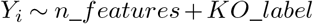

The coefficient of the knock-out label was tested using ANOVA. To perform differential expression of knock-out versus control in the multiome data, we performed a Wilcoxon rank-sum test in each Louvain cluster separately using the presto R package (version 1.0.0) (61). Multiple-testing correction was applied to all results using the Benjamini-Hochberg method to obtain FDR-adjusted p values.

#### Differential cell-cell communication analysis for the GLI3 knock-out experiment

To study how the cell-cell communication patterns were changed in the GLI3 KO organoids, CellPhoneDB (version 2)(53) with the default setting was applied to the single-cell gene expression in the single-cell multiome data of the three-week GLI3 KO and control organoids separately, given the integrated clustering results, to identify LR pairs significantly co-expressed in pairs of cell clusters (permutation test, P<0.05). For each of the source-target cluster pairs, the number of significant LR pairs was counted and compared between the two conditions. LR pairs with significant co-expression in at least one source-target cluster pair in both conditions were identified. COMUNET (54) was then applied to calculate expression pattern dissimilarities of the shared LR pairs between the two conditions. Shared LR pairs with dissimilarity > 0.8 were considered as changed LR pairs. Based on the shared LR pairs, ΔDegree was calculated for each cluster, as the difference between out-degree (summed strength of LR pairs with ligand secreted from the cluster) and in-degree (summed strength of LR pairs with receptors presented in the cluster).

#### Differential accessibility analysis for the GLI3 knock-out experiment

To find peaks with differential accessibility between GLI3 knock-out and control, we fit a generalized linear model with binomial noise and logit link for each peak i on binarized peak counts Y with the total number of fragments per cell and the knock-out label as the independent variables:

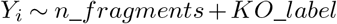

In addition, we fit a null model, where the knock-out label was omitted:

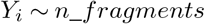

We then used a likelihood ratio test to compare the goodness of fit of the two models using the lmtest R package (version 0.9) (https://cran.r-project.org/web/packages/lmtest/index.html). Multiple testing correction was performed using the Benjamini-Hochberg method.

#### Comparison of perturbation effects with gene regulatory network

Before using the GRN to interpret the DE results, we first sought to assess the degree to which the transcriptomic effects of the GLI3 knock-out are consistent with the inferred GRN. We tested the enrichment of DE genes in the first (direct) and second order (indirect) neighborhood of GLI3 in the GRN graph using a fisher exact test. Further, we computed the shortest path from GLI3 to every DE gene in the GRN graph. To test how accurately the GRN can be used to predict the directionality of the DE, we computed the combined direction of each path as the product of the signs of all individual edges. We then determined the overall predicted effect of GLI3 on each DE gene by computing the mode of the directions of all shortest paths leading to that gene. We defined accuracy as the fraction of genes for which the DE direction was the inverse of the predicted overall effect. Next, we further filtered the paths so that all paths were composed only of DE genes and the direction of each path and subpath was consistent with the DE direction. To visualize this subgraph, we further pruned the graph by only retaining the path with the lowest average *log*_10_ p value for each DE gene.

#### Functional annotation of differentially accessible genomic regions

To better functionally assess the epigenomic effects of the GLI3 knock-out, we performed functional enrichment analysis with GREAT (35). We performed differential accessibility analysis in clusters 0 and 2 (early telencephalon) and applied a FDR threshold of 10^−4^. From all differentially expressed peaks, we selected the top 5000 peaks with lowest (most negative) linear model coefficient (depleted in the KO). We further selected all peaks that were accessible in at least 1% of cells in these clusters as the set of background peaks. Using these two peak sets, we used GREAT with the GRCh38 genome assembly and otherwise default parameters to obtain functional enrichment results. We reported GO Biological Process enrichments with FDR < 0.01 and that were supported by >100 foreground regions.

---

**Fig. S1.**
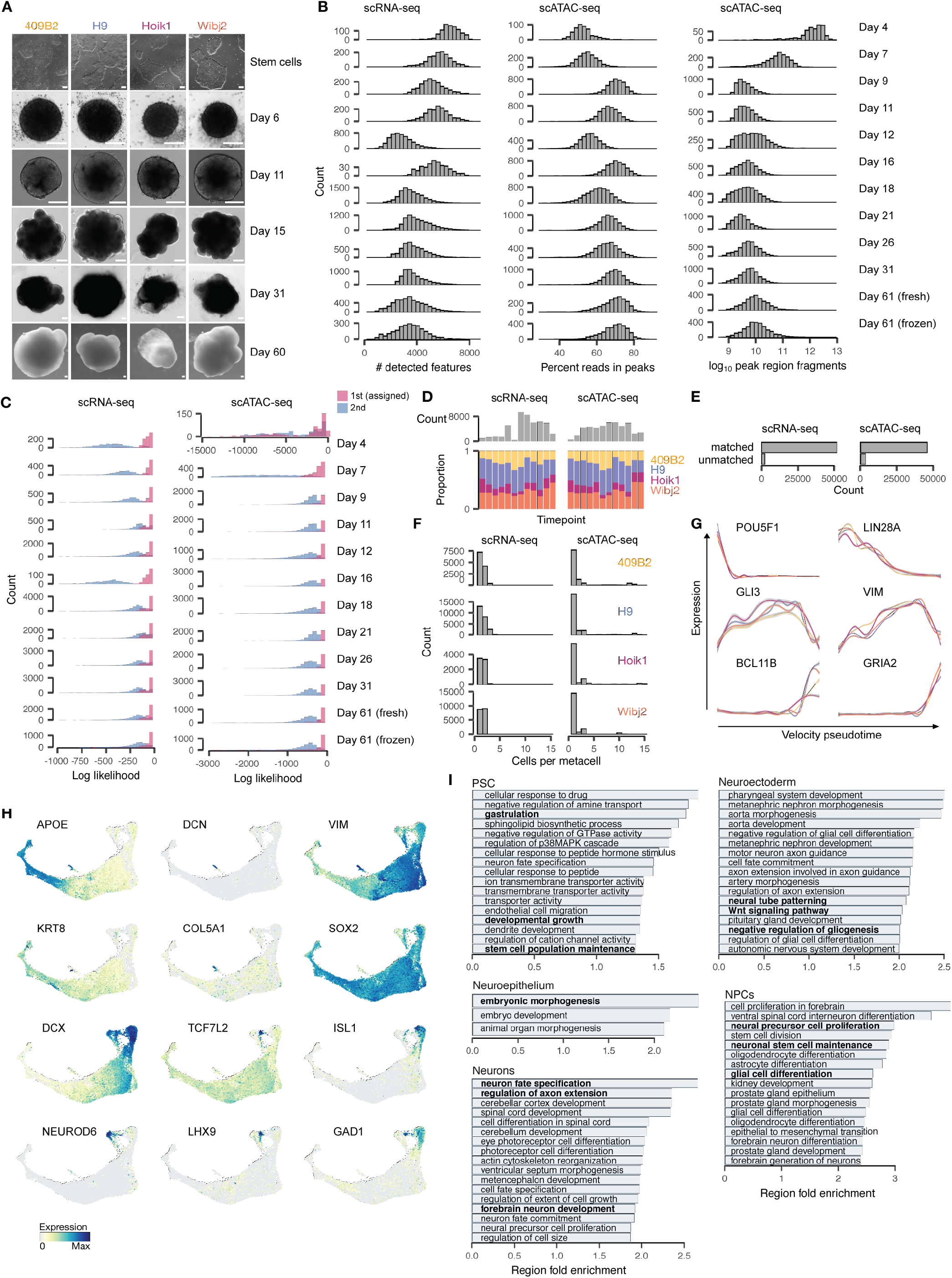
Supplemental analysis of cerebral organoid developmental multiome data. (A) Phase contrast (stem cells to day 15) and bright field (day 31 and day 60) showing examples of different stages of organoid development for four different stem cell lines. Scale bar is 200 µm. (B) Histogram of scRNA-seq and scATAC-seq quality control metrics for every time point. (C) Histograms showing counts of the first and second iPSC line assigned for each cell after demultiplexing based on single nucleotide variants. (D) Bar plot of number of cells for each timepoint (top) and stacked barplot showing proportion of cell lines (bottom) at different time points of scRNA-seq and scATAC-seq datasets. (E) Pseudo temporal expression patterns of stem cell (top), neuronal progenitor cell (middle) and neuronal (bottom) markers for each line. (F) Bar plots showing number of cells that were matched and unmatched with minimum-cost, maximum-flow (MCMF) bipartite matching in CCA space. (G) Histogram showing the number of cells that were matched per scRNA-seq/sc-ATACseq metacell for each cell line. (H) UMAP embedding colored by selected marker genes. (I) Barplots showing the results of the functional enrichment analysis of stage specific peak sets with GREAT. Significant (FDR < 0.01) GO Biological Process terms with highest fold enrichment are shown. Terms shown in Figure 1 are highlighted in bold.

**Fig. S2.**
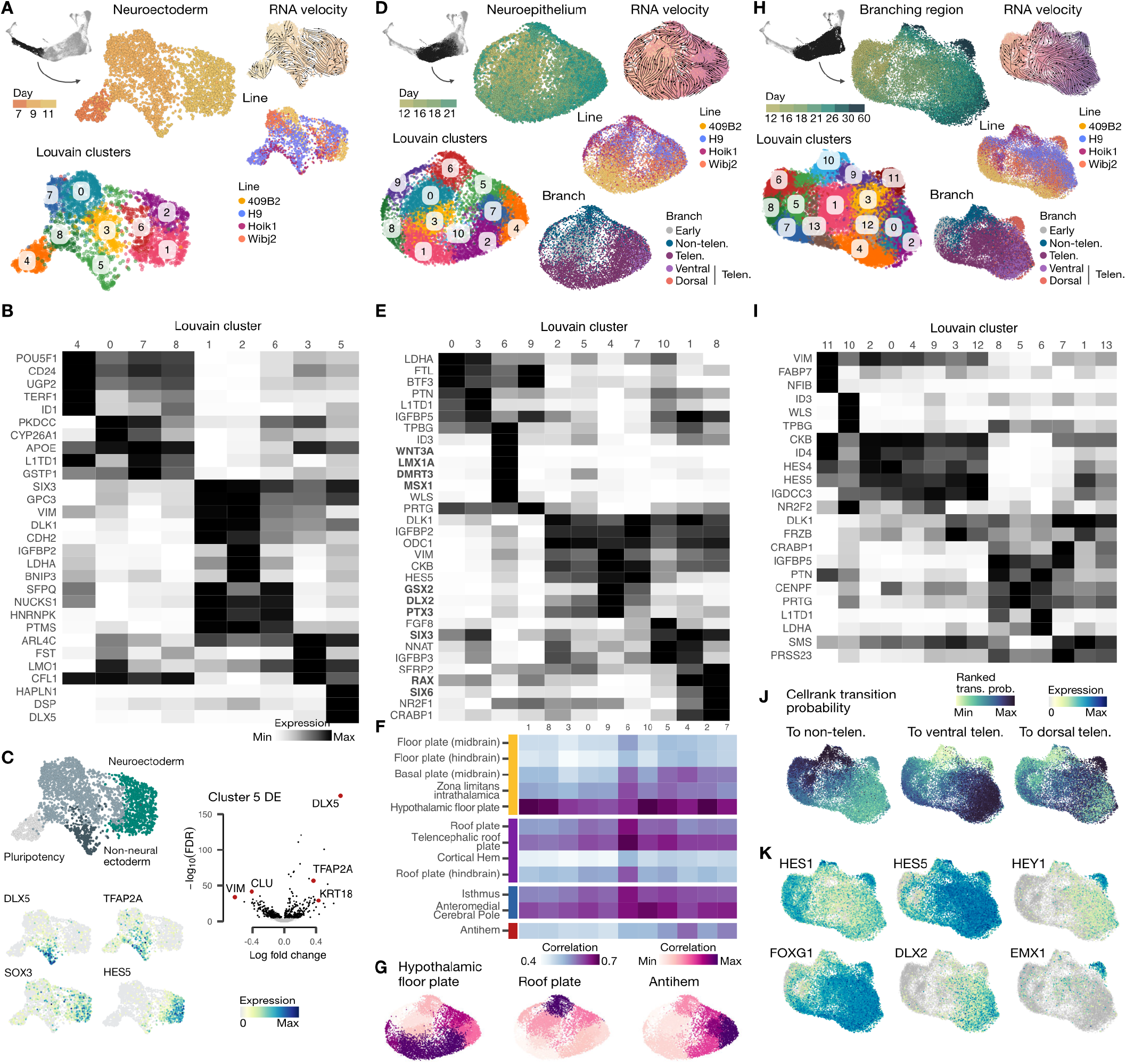
Heterogeneity analysis in different stages of organoid development. (A) UMAP embedding of a subset of the organoid trajectory surrounding neuroectoderm cells colored by time point, velocity pseudotime, cell line and Louvain clusters. (B) Heatmap showing averaged cluster expression of cluster markers. (C) UMAP embedding colored by cluster identities, expression patterns of cluster markers. Volcano plot shows differentially expressed (DE) genes of cluster 5 relative to other clusters. (D) UMAP embedding of a subset of the organoid trajectory surrounding neuroepithelial cells colored by time point, velocity pseudotime, cell line, branch prediction and lovain clusters. (E) Heatmap showing averaged cluster expression of cluster markers and manually selected marker genes of mouse developing brain organizer populations (24) (bold). (F) Heatmap showing correlation of different clusters to mouse developing brain organizers (24). (G) UMAP embedding colored by the correlation to developing mouse brain organizers. (H) UMAP embedding of a subset of the organoid trajectory colored by time point, velocity pseudotime, cell line, branch prediction and lovain clusters. (I) Heatmap showing averaged cluster expression of cluster markers. (J) UMAP embedding colored by rank-transformed CellRank transition probability to non-telencephalon, ventral telencephalon and dorsal telencephalon. (K) UMAP embedding colored expression of selected transcription factors.

**Fig. S3.**
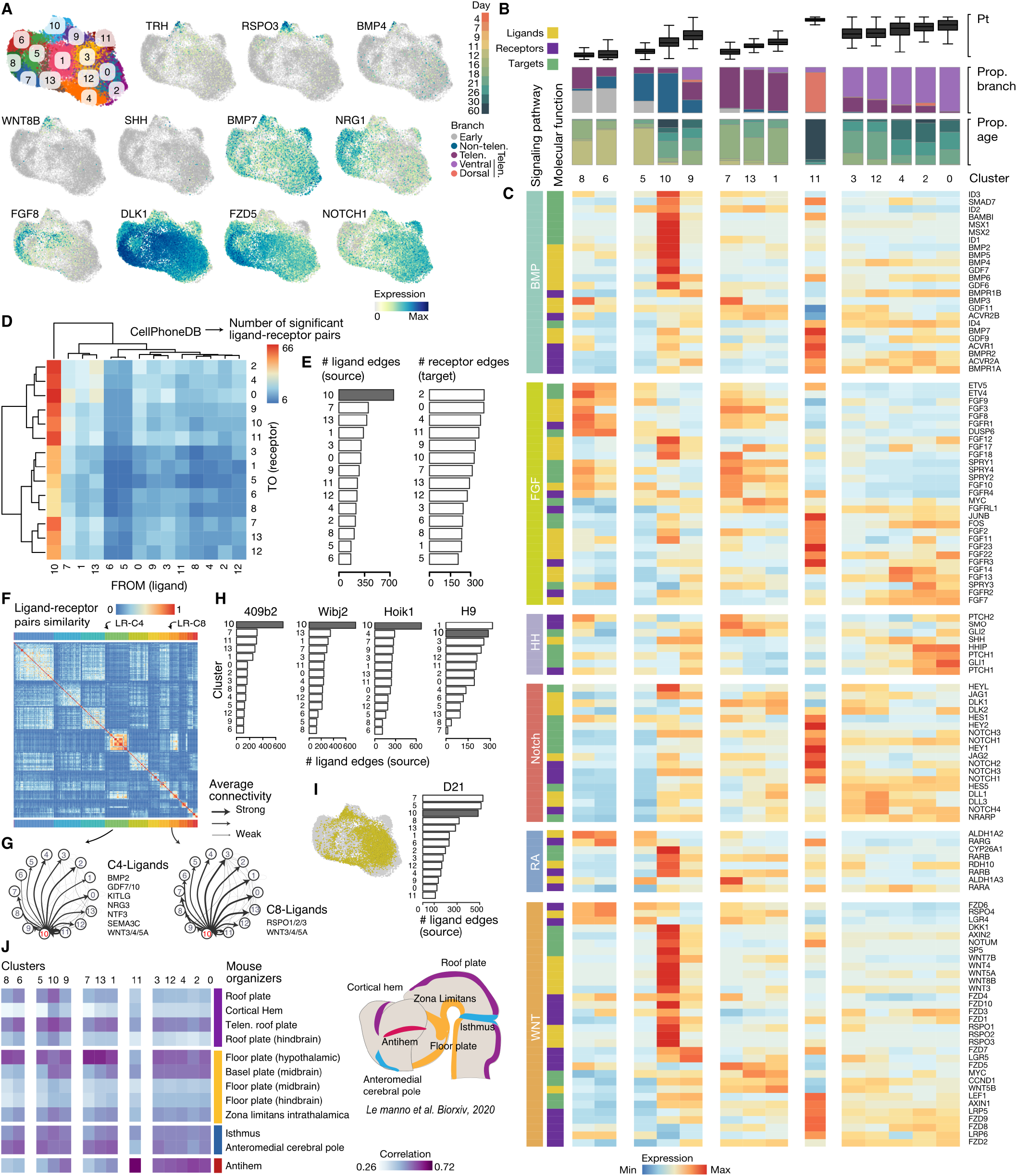
Receptor-ligand pairing analysis of cerebral organoid patterning. (A) UMAP embedding of a subset of neural progenitor cells (see Figure 2A) colored by Louvain clusters (top left) and normalized expression of signaling molecules. (B) Distribution of pseudotime position (top), proportion in each brain region (middle), and proportion at each time point (bottom) are shown for cells in each Louvain cluster. (C) Heatmap shows normalized expression of genes from different signaling pathways in each cluster, annotated as receptor, ligang, or transcription factor target. (D) Heatmap shows relative number of ligand (from) and receptor (to) pairs between each cluster identified using an interaction analysis (CellPhoneDB). (E) Barplot shows the number of edges from the interaction analysis that include a ligand (left) or receptor (right) for each cluster. (F) Heatmap showing similarities between different ligand-receptor pairs estimated by COMUNET, based on their inferred signaling strengths and directions between different cell clusters. (G) Barplot shows the number of edges from the interaction analysis that include a ligand for each cluster separated out by PSC line. (I) UMAP colored by cells at the day 21 time point (left) and barplot showing the number of ligand edges at this time point (right). (J) Heatmap showing correlation of different clusters to mouse developing brain organizers (left) with a schematic representation of the mouse neural tube marking the localization of organizers (right). Colors group early organizers by ventralizing (yellow), dorsalizing (purple) and boundary-inducing (cyan) activity. Late organizers are in pink. Adapted from (24).

**Fig. S4.**
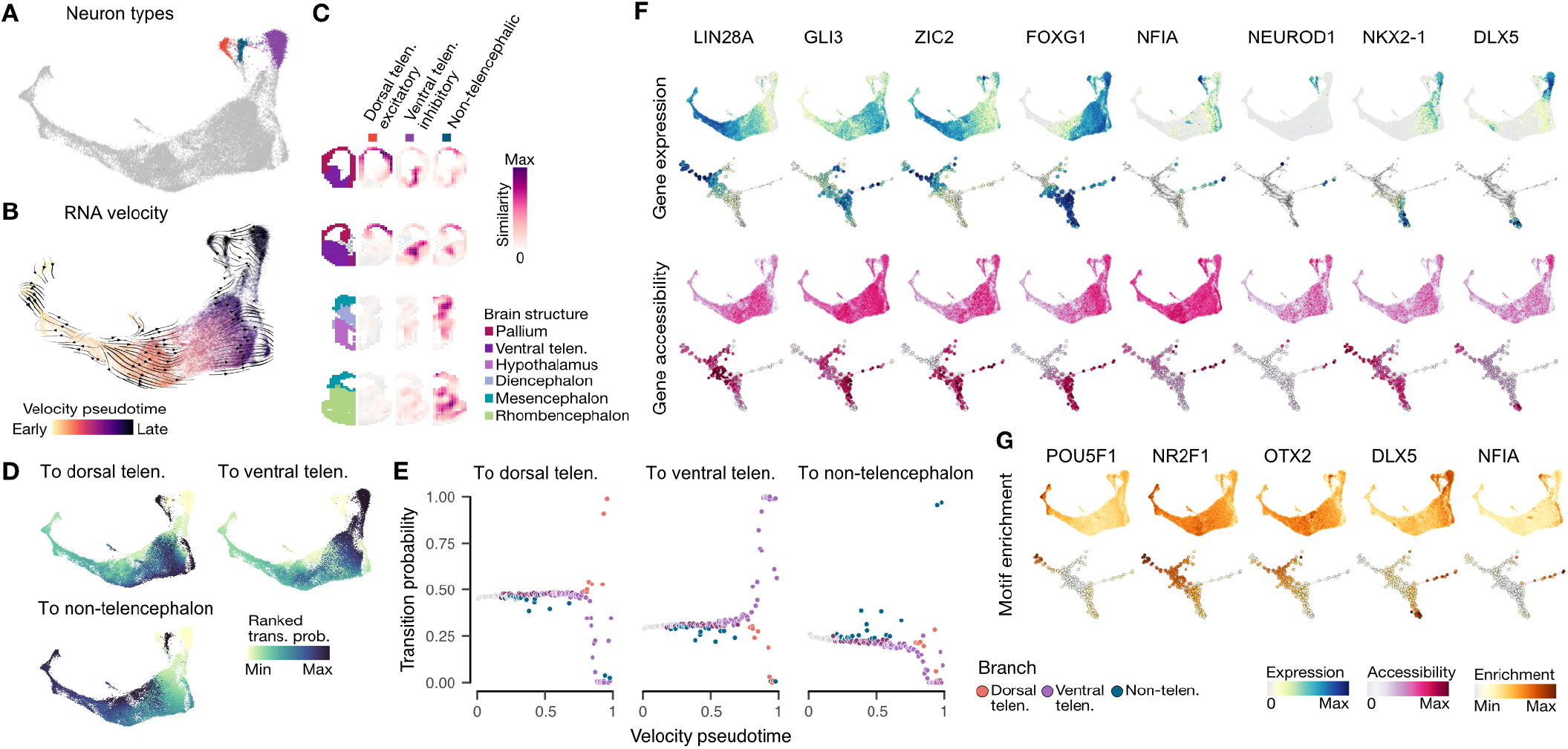
Trajectory reconstruction in the multi-omic developmental atlas. (A-B) Time course UMAP embedding colored by neuron types (A) or RNA velocity pseudotime (B). (C) Voxhunt plots showing expression similarity of neuron subtypes in cerebral organoids to voxels in five example sections of the developing mouse brain (embryonic day 13.5), as well as the structural annotation of the sections. (D) UMAP embedding colored by ranked transition probabilities. (E) Scatter plot showing transition probabilities as computed by CellRank versus velocity pseudotime. Each dot represents one high-resolution cluster. (F) UMAP embedding of the integrated time course and graph embedding colored by gene expression (top) and gene accessibility (bottom) for selected marker genes. (G) UMAP and graph representation colored by transcription factor motif enrichment for selected motifs.

**Fig. S5.**
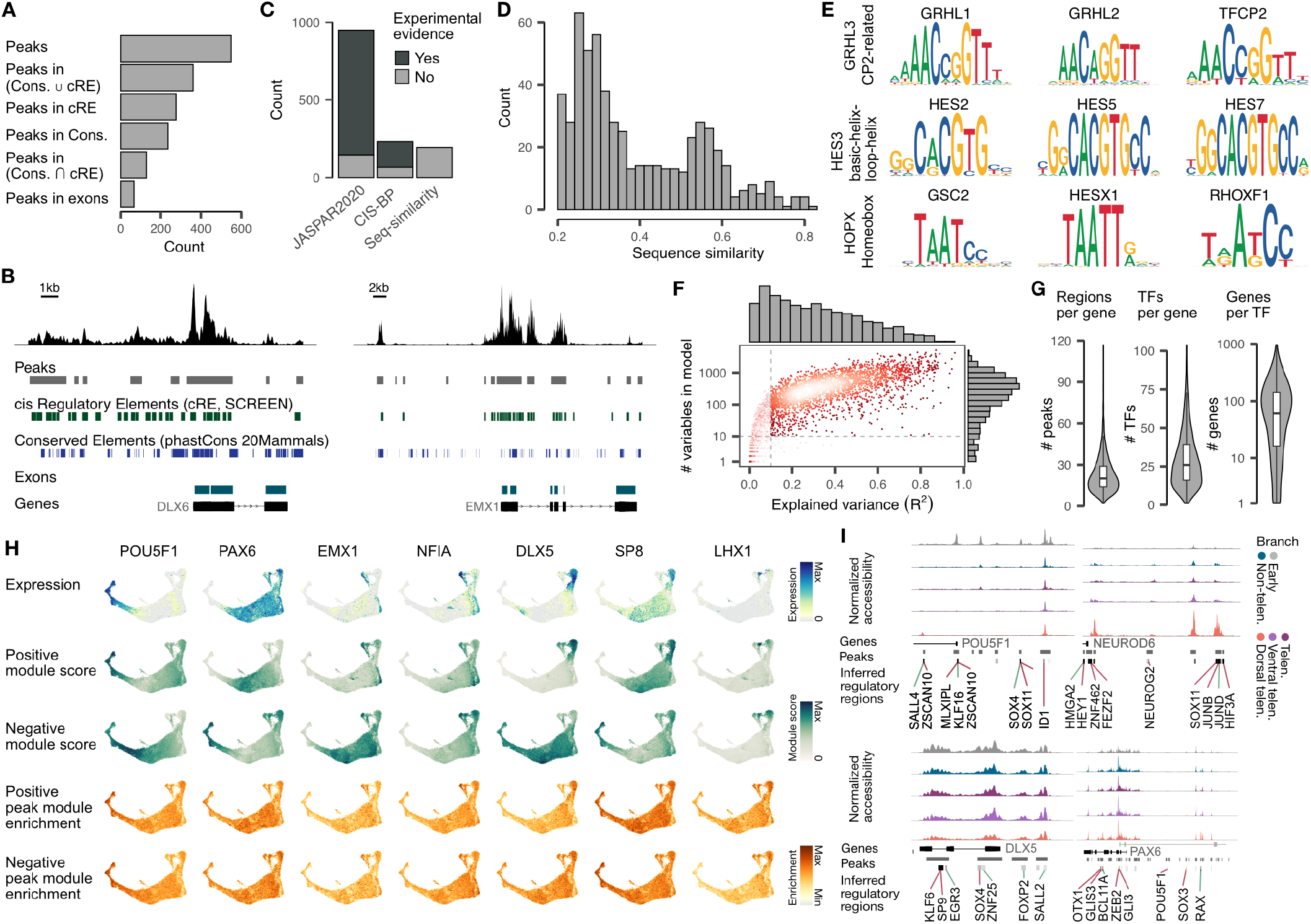
Gene regulatory network features of cerebral organoid development. (A) Numbers of chromatin access peaks in non-protein coding conserved regions (Cons.), previously annotated cis regulatory regions (cRE), or protein coding regions (exons). (B) Representative loci showing chromatin access (top) overlaying peak, cRE, conserved element coordinates, and exon coordinates. (C) Barplot showing the number of motifs used in GRN construction from two curated databases (JASPAR, CIS-BP), as well as motifs assigned through TF amino acid sequence similarity (Seq-similarity). (D) Histogram showing sequence similarity distribution of assigned TFs to query TF. (E) Examples of 3 TFs that have no motif annotation that were assigned motifs of other TFs based on amino acid sequence similarity. (F) Density scatter plot and histograms show the relationship between and distributions of explained variance (x) and number of variables (y) in the fitted models for GRN construction. (G) Violin plots show the distribution of peaks (left) and TFs assigned per gene (middle), and number of genes assigned per TF (right). (H) UMAP representation of time course colored by gene expression and module activity (rows) that can be extracted from the GRN for representative TFs (columns). (I) Representative loci showing average chromatin access signal tracks at different branches (PSC/Neuroectoderm, grey; Telencephalon progenitors, dark purple; Dorsal telencephalon, plum; Ventral telencephalon, purple) overlaying inferred transcription factor binding sites within regulatory regions.

**Fig. S6.**
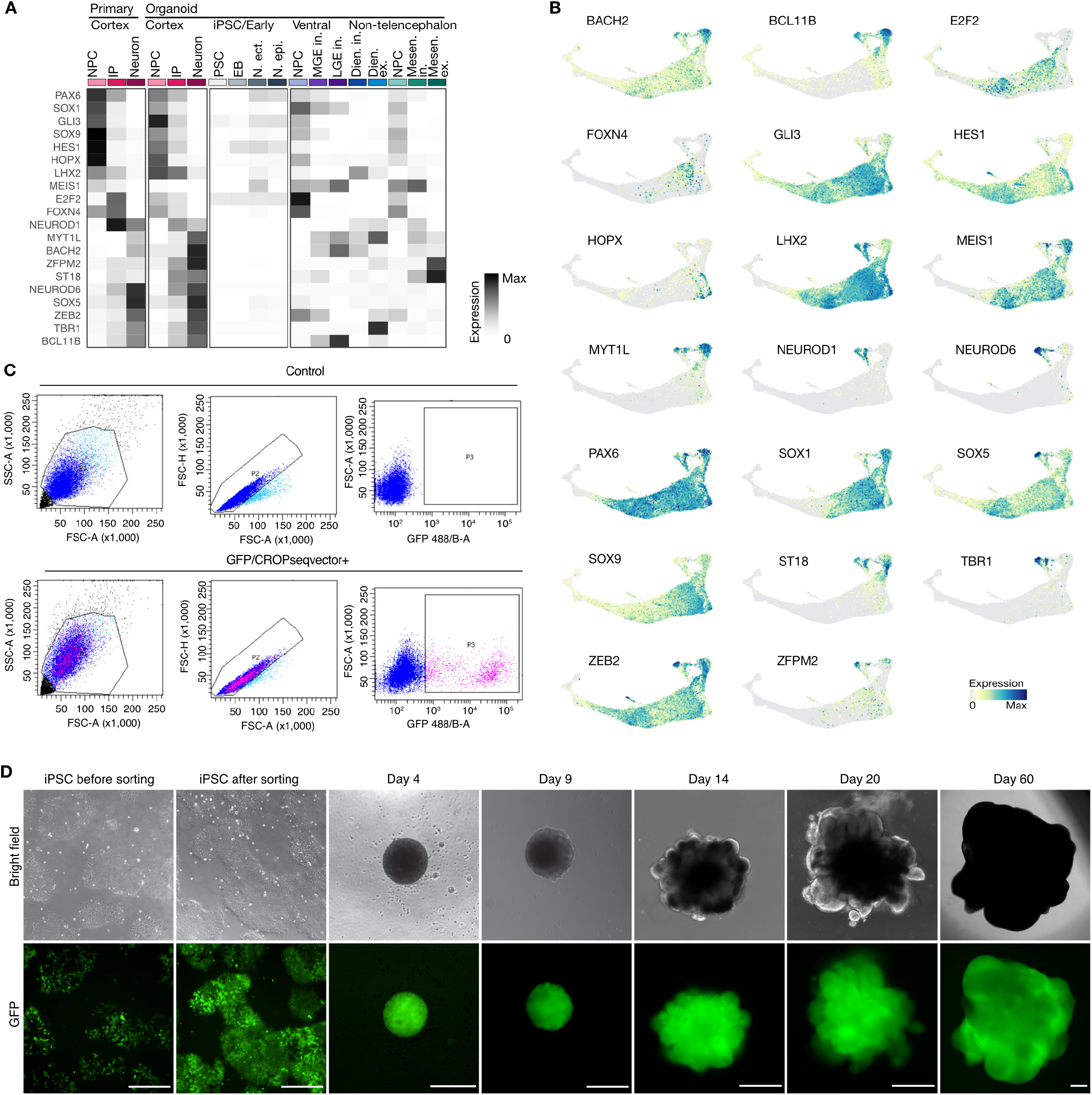
Target selection and experimental details for the single-cell in organoid perturbation experiment. (A) Averaged expression of genes targeted in the single-cell genomic perturbation experiment in neuronal progenitors (NP), intermediate progenitors (IP) and neurons in the primary human and organoid developing cortex, as well as in induced pluripotent stem cells (IPSC), the embryoid body (EB), ventral telencephalic NPCs, inhibitory neurons of the medial ganglionic eminence (MGE in.), lateral ganglionic eminence (LGE in.), non-telencephalic NPCs, diencephalic excitatory neurons (Dien. ex.) and inhibitory neurons (Dien. in.) and mesencephalic excitatory neurons (Mesen. ex.) and inhibitory neurons (Mesen. in.). (B) UMAP embedding colored by the expression of all targeted genes. (C) Exemplary Fluorescence-activated cell sorting plots of the sorting scheme used to isolate CROP-seq vector positive induced pluripotent stem cells (iPSCs). (D) Phase contrast and CROP-seq vector positive (GFP) imaging during cerebral organoid development. Scale Bar is 500 µm.

**Fig. S7.**
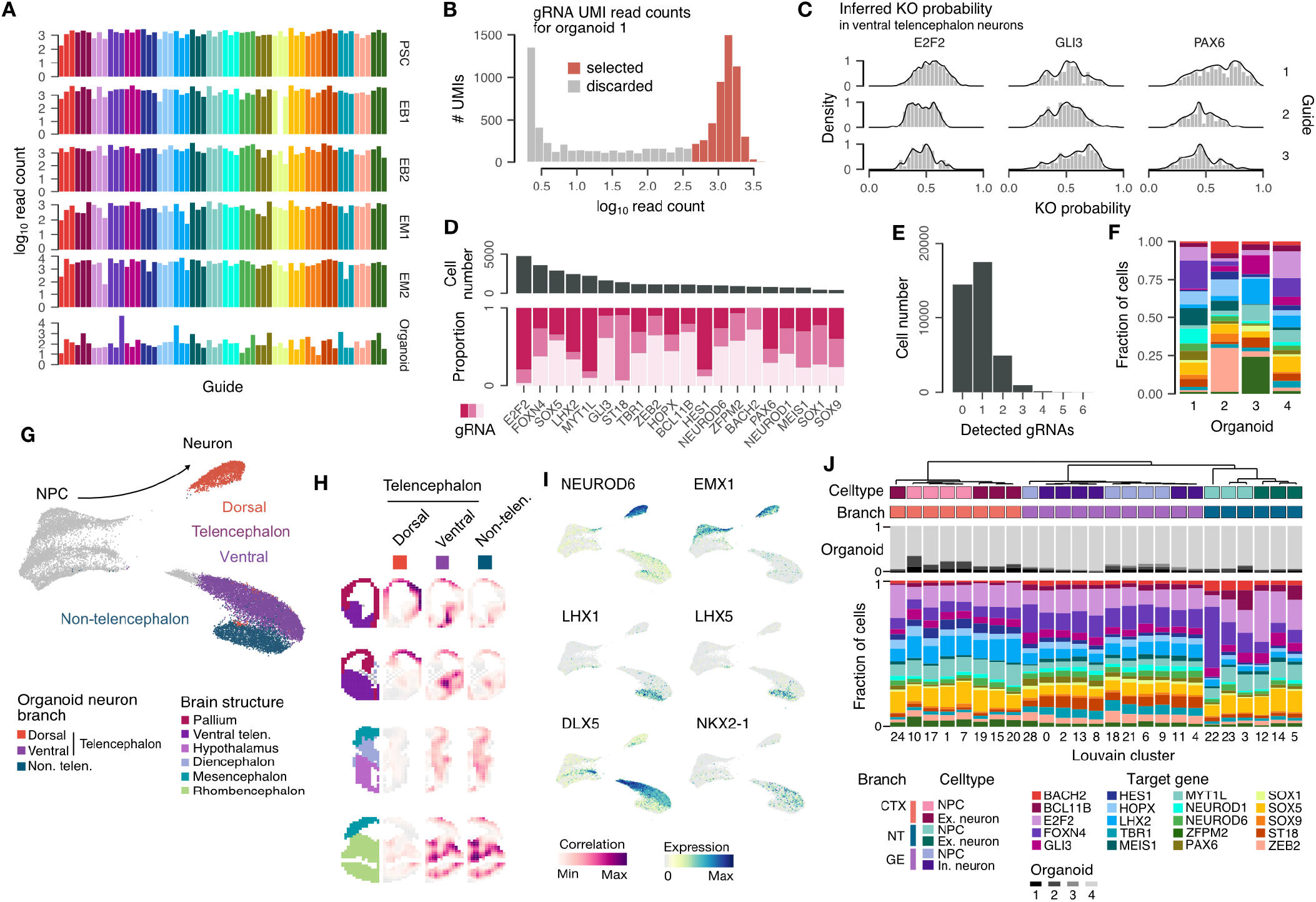
Guide detection and celltype annotation in the single-cell in organoid perturbation experiment. (A) Barplot showing number of cells with detected guide RNA (gRNA) for each targeted gene and stacked barplot showing the distribution of the different gRNAs targeting the same gene. (B) Histogram showing the distribution of read counts for gRNA UMIs after amplicon sequencing for one organoid. UMIs marked in red were selected for downstream analyses. (C) Density histograms showing the distribution of inferred KO probabilities for gRNAs of 3 different target genes. (D) Barplot showing cell number and proportion of gRNAs for all target genes. (E) Barplot showing the number of guides detected in sequenced cells. (F) Barplot showing the proportion of cells with each perturbation for all organoids (G) UMAP embedding colored by annotated neuron subtypes. (H) Voxhunt plots showing expression similarity of neuron subtypes in cerebral organoids to voxels in five example sections of the developing mouse brain (embryonic day 13.5), as well as the structural annotation of the sections (left). (I) UMAP embedding colored by expression of non-telencephalic (top), ventral (middle) and dorsal (bottom) neuron markers. (J) Hierarchical clustering of Louvain clusters based on the composition of gRNAs targeting different genes. Cell type and branch annotations are shown as side bars. Compositions of organoids and composition of cells with gRNAs targeting different genes are shown below as stacked bar plots.

**Fig. S8.**
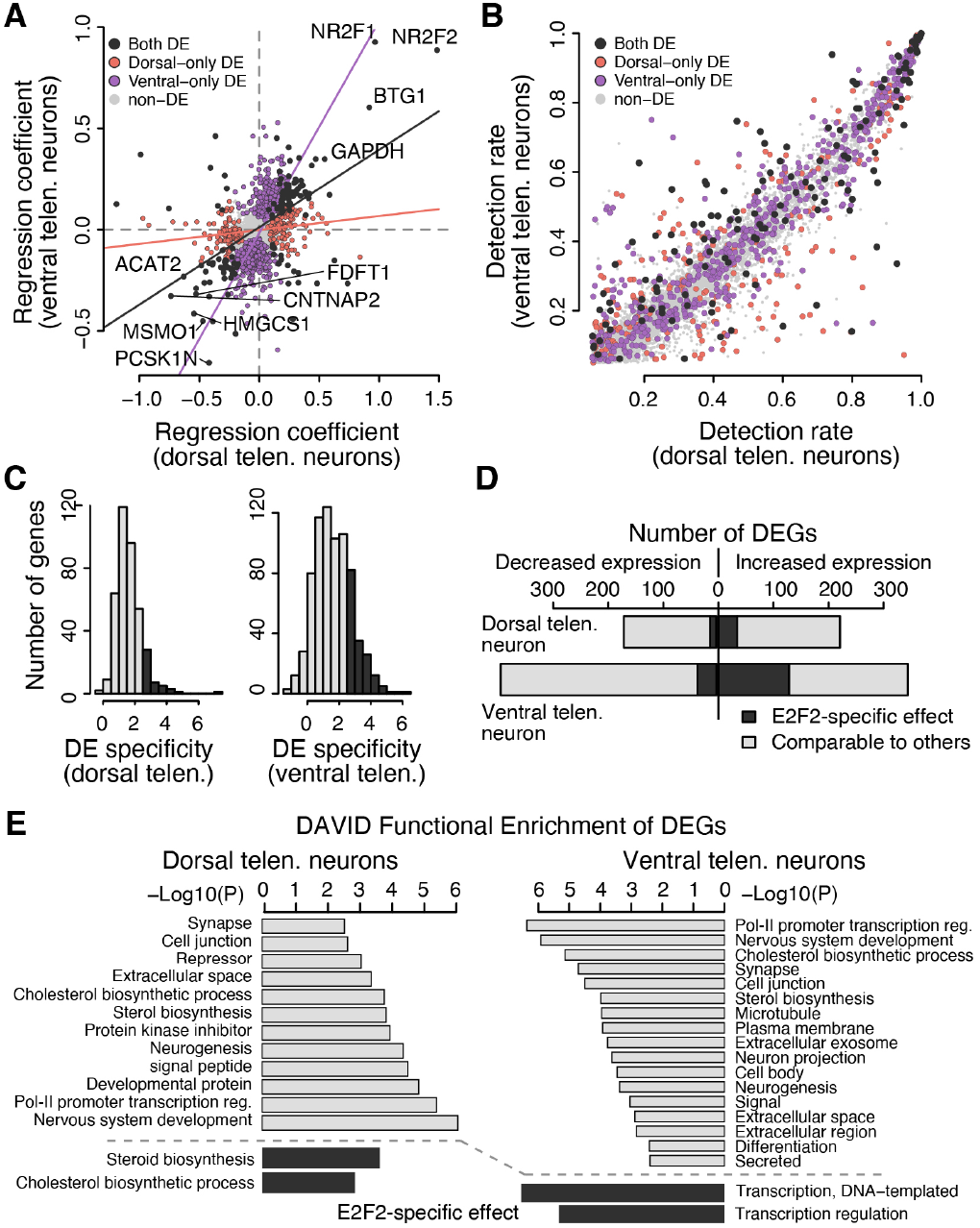
Differential expression of dorsal and ventral telencephalic neurons with E2F2 targeting gRNAs. (A) Scatter plot shows expression changes between neurons with E2F2 targeting gRNAs and other neurons in dorsal (x-axis) and ventral (y-axis) telencephalic neurons, with each dot representing one gene. Colors of dots represent the neuron types where differential expression is detected. Lines show the correlation of expression changes in the two neuron types, with DE genes in both types and DE genes in only one type shown separately. (B) Scatter plot shows detection rates of genes in dorsal (x-axis) and ventral (y-axis) telencephalic neurons, colored by the neuron types where differential expression is detected. (C) DE specificity of the identified E2F2 DEGs in dorsal (left) and ventral (right) telencephalic neurons, relative to the differential expression detected in cells with other targeting gRNAs. Dark bars show DE genes with specificity > 2.5 and are considered to show E2F2-specific effect. (D) Stacked bar plot shows numbers of DE genes with increased or decreasing in the two neuron types. The dark bars show DEGs with E2F2-specific effects. (E) Examples of functional enrichment for E2F2 DEGs in dorsal and ventral neurons with DAVID. Grey bars show enriched terms of all E2F2 DEGs, and dark bars show enriched terms of DEGs with E2F2-specific effects.

**Fig. S9.**
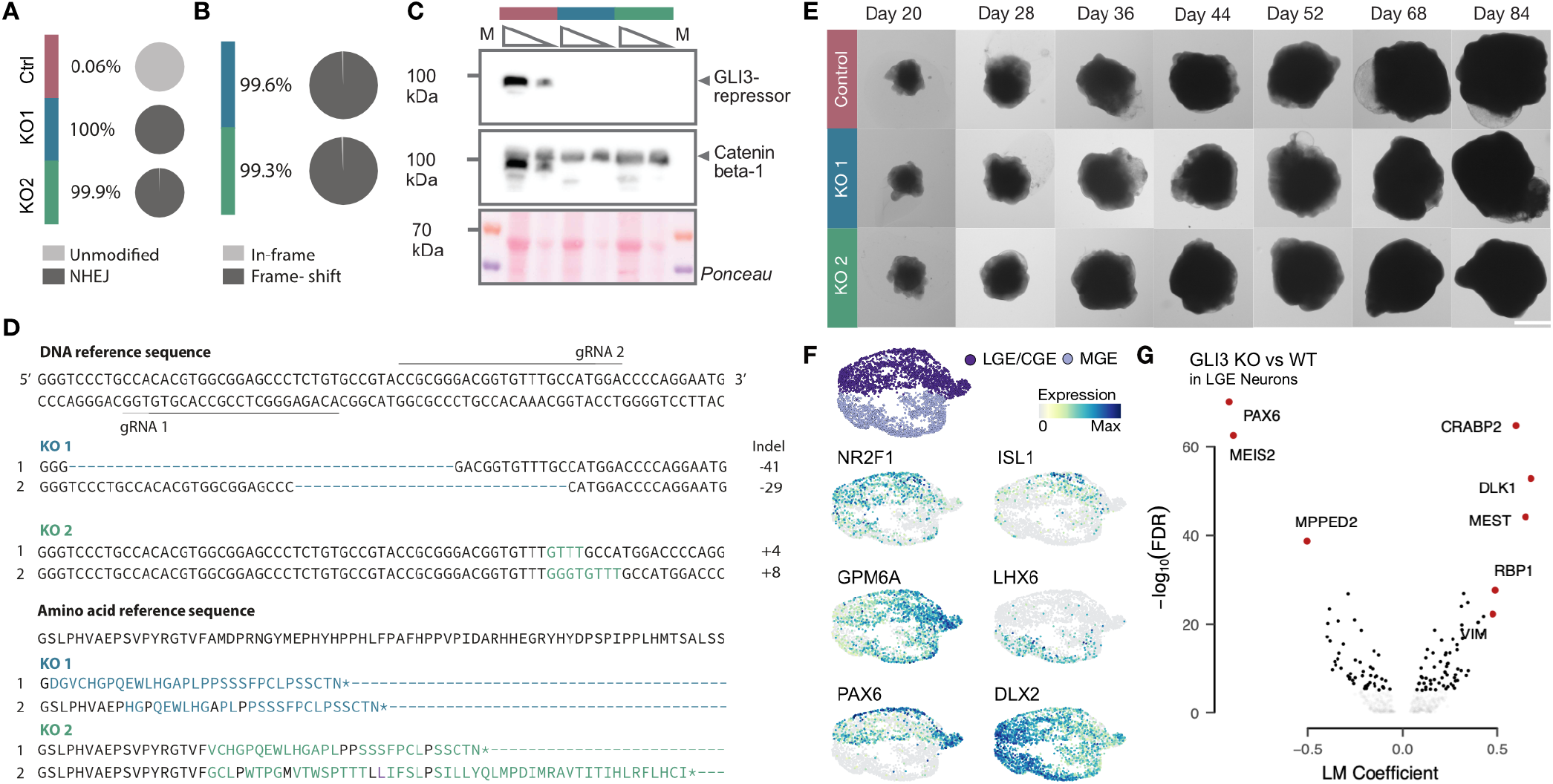
Characterization of GLI3 knock-out organoids. (A) Quantification of editing frequency as determined by the percentage and number of sequencing reads showing unmodified and modified alleles for the control and both KO cell lines. (B) Frequency of frameshift of coding sequence reads as a result of the modifications seen in both KO lines. (C) Western blot showing expression of Gli3-repressor (83kDA) in the control cell line. Catenin beta-1 and Ponceau were used a s loading control. (D) Sequences of the coding strand of the different indels of the different KO lines. The reference sequence is corresponding with the control line. The position of the gRNAs with the protospacer adjacent motif (PAM)-sequence is depicted above and underneath the sequence. Reference protein sequence with the protein sequences of each KO line of the altered protein sequences caused by the frame-shift. (E) Brightfield images of cerebral organoid development with control and both KO cell lines. Scale bar is 2 mm. (F) UMAP embedding of ventral telencephalic GLI3 KO neurons showing medial ganglionic eminence (MGE) and lateral/caudal ganglionic eminence (LGE/CGE) neuronal populations (top). Feature plots show selected marker gene expression on the UMAP embedding. (G) Volcano plot showing differential expression analysis in LGE neurons for GLI3 WT versus KO cells.

**Fig. S10.**
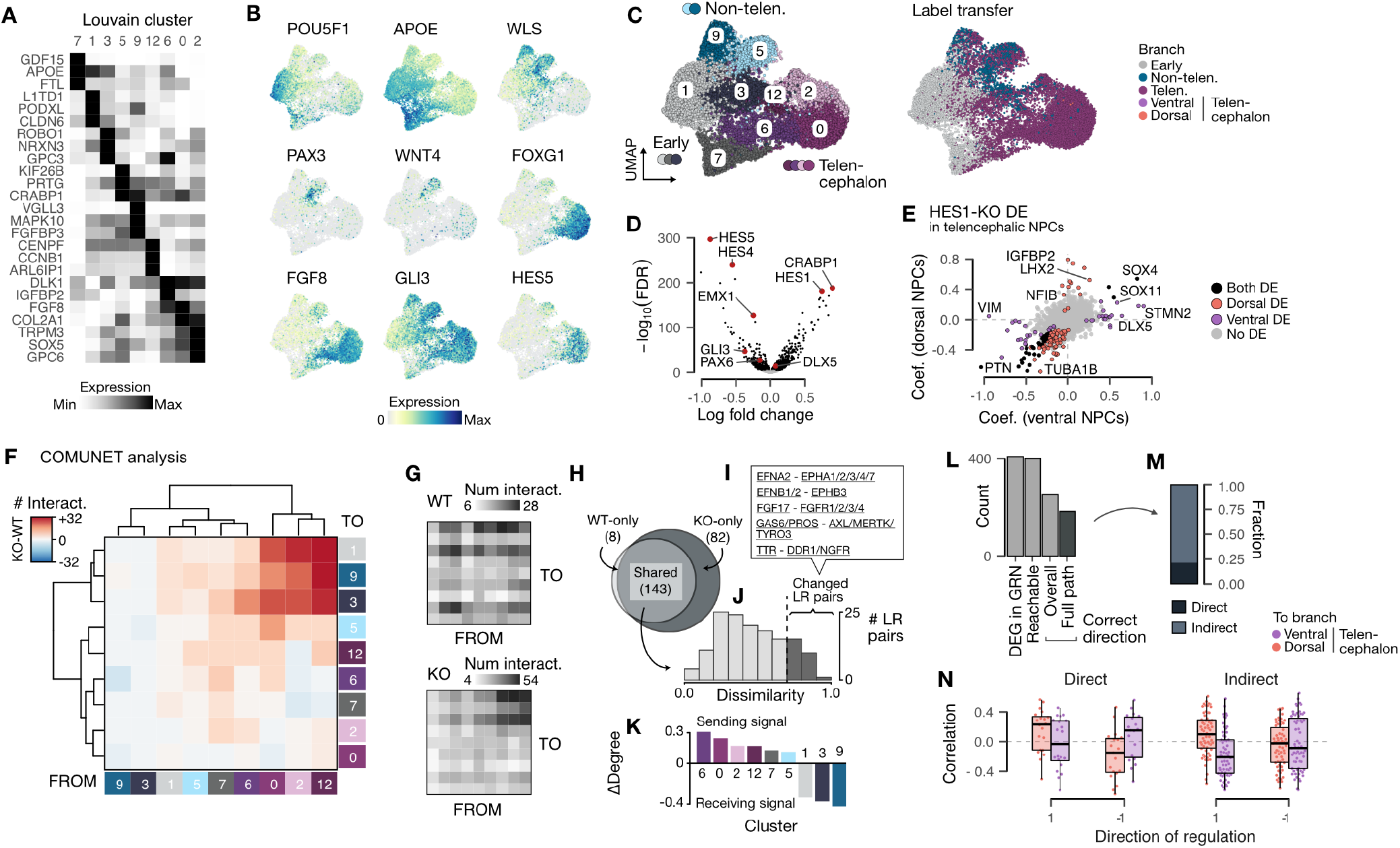
GLI3 KO induced changes in telencephalic progenitors in cerebral organoids. (A) Heatmap showing the expression of top marker genes for unbiased Louvain clusters. (B) UMAP colored by the expression of selected genes marking pluripotency (POU5F1, APOE), early non-telencephalic progenitors (WLS, PAX3, WNT4) and early telencephalic progenitors (FOXG1, FGF8, GLI3, HES5). (C) UMAP embedding colored by Louvain clusters and by regional branch labels predicted by label transfer from the organoid developmental time course atlas. (D) Volcano plot showing the differential expression of genes in telencephalic progenitors upon GLI3 KO. (E) Scatter plot showing differential gene expression in neural progenitor cells (NPCs). (F) Heatmap showing the changed numbers of ligand (from) and receptor (to) pairs between each cluster identified using an interaction analysis (CellPhoneDB) between GLI3 KO and control cells. (G) Heatmap showing the number of ligand (from) and receptor (to) pairs between each cluster in control (upper) and GLI KO (lower) cells. (H) Venn diagram showing number of identified LR pairs in each of the two conditions as well as their shared ones. (I) Histogram showing the distribution of COMUNET-based dissimilarities of the shared LR pairs between the control and GLI3 KO conditions. (J) Examples of the changed LR pairs between the two conditions. (K) Bar plot showing ΔDegree of each cluster, calculated based on the changed LR pairs. Positive ΔDegrees represent clusters sending signals, and negative ones represent clusters receiving signals. (L) Barplot showing the number of all DE genes present in the GRN (DEG in GRN), all DE genes that are reachable from GLI3 in the GRN graph (Reachable), DE genes where the overall direction predicted by the GRN was consistent with the DE result (Overall) and DE genes for which all subpaths on the shortest path from GLI3 were individually consistent with the DE result (Full path). (M) Barplot showing the fraction of DE genes directly and indirectly regulated by GLI3. (N) Boxplot showing the spearman correlation of the expression of directly and indirectly regulated DE genes with transition probabilities to dorsal telencephalon (top) and ventral telencephalon neurons (bottom). The x-axis indicates the inferred direction of regulation from GLI3, which is consistent with the DE.

## Notes

https://doi.org/10.5281/zenodo.5242913

https://github.com/quadbiolab/organoid_regulomes

https://github.com/quadbiolab/Pando

### Competing Interest Statement

The authors have declared no competing interest.

