## Supplementary figures and images for "Inferring and perturbing cell fate regulomes in human cerebral organoids"

### Data_S8.pdf

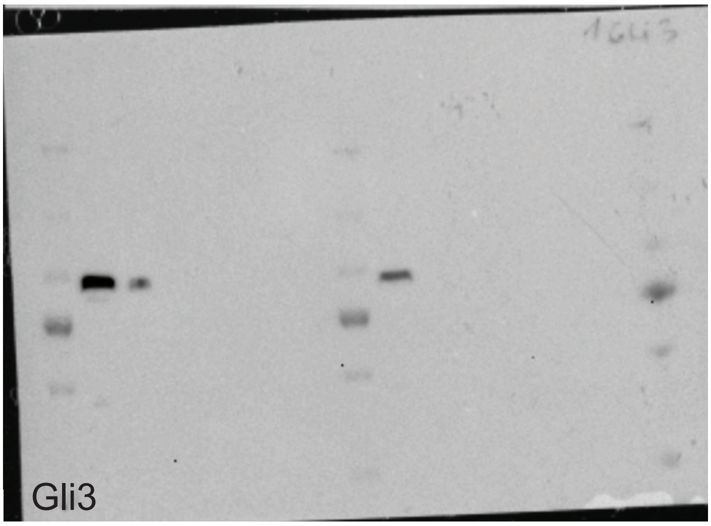


250 kDa
130 kDa
100 kDa
70 kDa
55 kDa
35 kDa

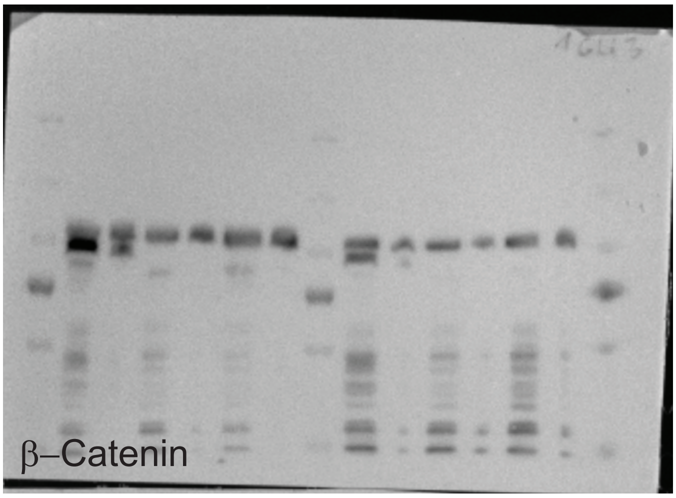


250 kDa
130 kDa
100 kDa
70 kDa
55 kDa
35 kDa

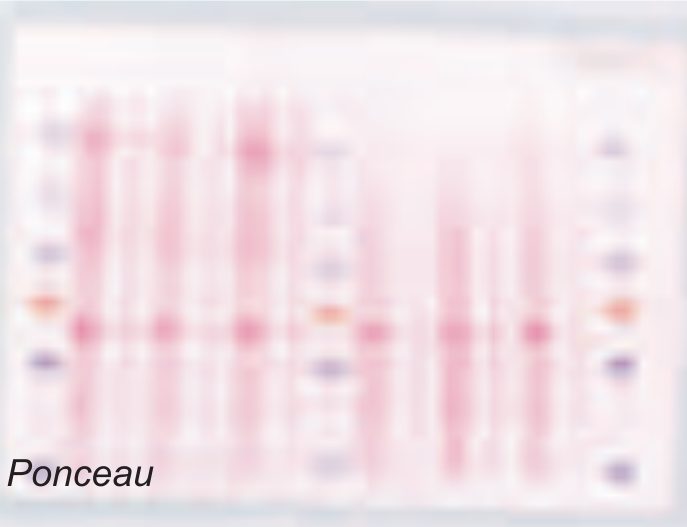


250 kDa
130 kDa
100 kDa
70 kDa
55 kDa
35 kDa
